# Oncogenic RAS Induces a Distinctive Form of Non-Canonical Autophagy Mediated by the P38-ULK1-PI4KB Axis

**DOI:** 10.1101/2024.12.10.627736

**Authors:** Xiaojuan Wang, Shulin Li, Shiyin Lin, Yaping Han, Tong Zhan, Zhiying Huang, Juanjuan Wang, Ying Li, Haiteng Deng, Min Zhang, Du Feng, Liang Ge

## Abstract

Cancer cells with RAS mutations exhibit enhanced autophagy, essential for their proliferation and survival, making it a potential target for therapeutic intervention. However, the regulatory differences between RAS-induced autophagy and physiological autophagy remain poorly understood, complicating the development of cancer-specific anti-autophagy treatments. In this study, we identified a form of non-canonical autophagy induced by oncogenic KRAS expression, termed RAS-induced non-canonical autophagy via ATG8ylation (RINCAA). RINCAA involves distinct autophagic factors compared to those in starvation-induced autophagy and incorporates non-autophagic components, resulting in the formation of non-canonical autophagosomes with multivesicular/multilaminar structures labeled by ATG8 family proteins (e.g., LC3 and GABARAP). We have designated these structures as RAS-induced multivesicular/multilaminar bodies of ATG8ylation (RIMMBA). A notable feature of RINCAA is the substitution of the class III PI3K in canonical autophagy for PI4KB. We identified a regulatory P38-ULK1-PI4KB-WIPI2 signaling cascade governing this process, where ULK1 phosphorylation at S317, S479, S556, and S758 activates PI4KB. This activation involves PI4KB phosphorylation at S256 and T263, initiating PI4P production, ATG8ylation, and non-canonical autophagy. Importantly, elevated PI4KB phosphorylation at S256 and T263 was observed in RAS-mutated cancer cells and colorectal cancer specimens. Inhibition of PI4KB S256 and T263 phosphorylation led to a reduction in RINCAA activity and tumor growth in both xenograft and KPC models of pancreatic cancer, suggesting that ULK1-mediated PI4KB-Peptide-1 phosphorylation could represent a promising therapeutic target for RAS-mutated cancers.

## INTRODUCTION

Macroautophagy (henceforth, autophagy) is the massive degradation of intracellular materials and is crucial for maintaining cellular homeostasis and survival under stress conditions.^1, 2^ During autophagy, damaged organelles, invasive bacteria, and aggregate-prone proteins are delivered to lysosomes for degradation.^1, 3^ Autophagy is relevant in the pathogenesis of diverse diseases, e.g., dysregulated autophagy is associated with cancer.^4–6^ However, the specific molecular events accounting for the dysregulation of autophagy, particularly the differences between cancer-associated autophagy and physiological autophagy, e.g., starvation-induced autophagy, has not been clarified.^7^

The *ras* family of genes, including *hras*, *kras*, and *nras* encode extremely similar 188– 189 amino acid proteins and are the most common oncogenes in human cancer.^8^ The *ras* genes encode monomeric GTPases, which function as molecular switches in signal transduction pathways regulating cell proliferation, differentiation, and survival of mammalian cells.^9^ Mutations that constitutively activate RAS proteins occur in 20–25% of all human cancers, with a *kras* mutation occurring the most frequently and in multiple cancer types.^10, 11^ Activation of a *kras* mutation is a key driver of cancer initiation and progression in many tumors and is essential for tumor growth.^12, 13^, However, hyperactive RAS mutants are difficult to target because of the lack of drug-binding pockets on the surface of the RAS proteins.^14^ Until now, the only effective inhibitor for clinical use was AMG510& MRTX849, which specifically targets KRAS (G12C).^15, 16^

In addition to directly targeting the RAS proteins, identifying drugs that inhibit downstream RAS effector proteins is a solution for treating cancers with RAS mutations.^17^ It has been shown that activating a RAS mutation associates with the overactivation of autophagy.^17–19^ Studies indicate that RAS-activated autophagy harbors unique features, such as providing nutrients for tumor growth, immune evasion, and remodeling the proteome via selective degradation, e.g., elimination of deleterious inflammatory response pathway components to prevent cytokine-induced paracrine cell death.^20–24^ Accordingly, RAS-mutant tumor cells are very sensitive to autophagy inhibitors, suggesting that autophagy can be exploited as a therapeutic target for RAS-mutant tumors.^25–27^ Because autophagy also maintains cellular homeostasis under physiological conditions, an autophagy inhibitor specifically preventing RAS-induced autophagy without affecting physiological autophagy should be developed. Nonetheless, the regulatory differences between RAS-mutation-induced autophagy and physiological autophagy are unclear, particularly regarding the different involvement of the core autophagy factors, i.e. autophagy-related genes (ATGs) and proteins.

The core steps in autophagy, including biogenesis and maturation of the autophagosome, are regulated by ATGs and other autophagy-related proteins.^28, 29^ Under starvation-induced autophagy (a physiological form of autophagy), components of the uncoordinated (UNC)- 51-like kinase (ULK) complex, including ULK1/2, ATG13, and RB1 inducible coiled-coil 1 (RB1CC1/FIP200), and ATG101 are activated by inhibiting the mechanistic target of rapamycin complex 1 (mTORC1) and meanwhile activating AMPK.^28, 29^ The ULK kinase can phosphorylate multiple autophagy regulators and one major target is the class III phosphoinositide 3 kinase (PI3K) complex comprised of Beclin-1, ATG14, phosphatidylinositol 3-kinase (PIK3) catalytic subunit 3 (PIK3C3/VPS34), PIK3 regulatory subunit 4 (PIK3R4/P150), and activating molecule in Beclin-1-regulated autophagy protein 1 (Ambra1).^28, 29^ The PI3K complex phosphorylates phosphatidylinositol (PI) lipids of the phagophore to recruit phosphatidylinositol-3- phosphate (PI3P)-interacting proteins, such as WD repeat domain phosphoinositide-interacting 2 (WIPI2). WIPI2 is an effector of PI3P that activates downstream events by recruiting the lipidation machinery (ATG5-ATG12 conjugate in complex with ATG16L1, ATG3, and ATG7). The lipidation machinery catalyzes the conjugation of microtubule-associated protein 1 light chain 3 (MAP1LC3/LC3/ATG8) to phosphatidylethanolamine (PE), which builds the autophagosome.^28, 29^ ATG9A seeds the autophagosome and acts as a scramblase together with ATG2, which transfers lipids to the autophagic membrane.^28, 29^ The ESCRT (endosomal sorting complex required for transport) complex seals the phagophore completing autophagosome biogenesis,^30^ and multiple SNARE complexes together with membrane tether components facilitate autophagosome to lysosome fusion.^29, 31, 32^ While the cascade and function of ATGs in physiological autophagy are well understood, the regulatory mechanisms governing these autophagic factors under pathological conditions remain less clear, particularly regarding their molecular distinctions from physiological autophagy.

Here, we employed multiple oncogenic RAS-induced autophagy models to investigate the underlying molecular mechanisms of autophagy dysregulation in cancer. Our results revealed that KRAS(G12V)-induced a non-canonical autophagy featured by ATG8ylation which operates independently of several autophagic factors typically associated with starvation-induced autophagy, such as class III PI3K, ATG9A and ATG2. We term it as RAS-induced non-canonical autophagy via ATG8ylation (RINCAA). Instead of double-membrane autophagosomes, alternative autophagosomes with multivesicular and multilaminar structures positive for ATG8 homologs (e.g LC3 and GABARAP) were observed. Specifically focusing on the PI3K independence, we discovered that activation of PI4KB by ULK1, induced by the RAS signal, is crucial. Notably, phosphorylation of Peptide 1on S256 and T263 by RAS-activated ULK1 enhances PI4KB activity, leading to PI4P production. Additionally, similar to starvation-induced autophagy, WIPI2 functions as a downstream effector but acts as a PI4P effector, instead of PI3P, initiating downstream events of autophagosome biogenesis. Inhibition of MEK1/2 and PI4KB phosphorylation on S256 and T263 in combination effectively suppressed autophagy and inhibited the proliferation of mutant-RAS tumor cell lines, and tumor growth in xenograft and KPC pancreatic cancer models. These findings elucidate a mechanistic framework for an oncogenic KRAS-induced alternative autophagy and shed lights on a new potent target against cancer with Ras mutations.

## RESULTS

### Establishing a cellular system to analyze KRAS-induced autophagy

The expression of a constitutively activated RAS mutant triggers autophagy. ^25, 33–35^ We were curious to know if RAS-induced autophagy employs the same cascade of ATGs as starvation-induced autophagy. Therefore, we engineered 293T cells with doxycycline (Dox)-inducible G12V-mutant KRAS gene (KRAS(G12V)) to activate autophagy. KRAS(G12V) expression was dose-dependent and, for subsequent autophagy analysis, we systematically adjusted the concentration and fine-tuned the induction duration to achieve an expression level comparable to that of endogenous RAS in various cancer cell lines. (Fig. 1a, b). We examined lipidated LC3 (LC3-II), an autophagic membrane marker. Western blot analysis showed that the Dox treatment increased LC3-II levels in the presence and absence of the lysosome inhibitor Bafilomycin A1, which blocks

**Fig. 1.**
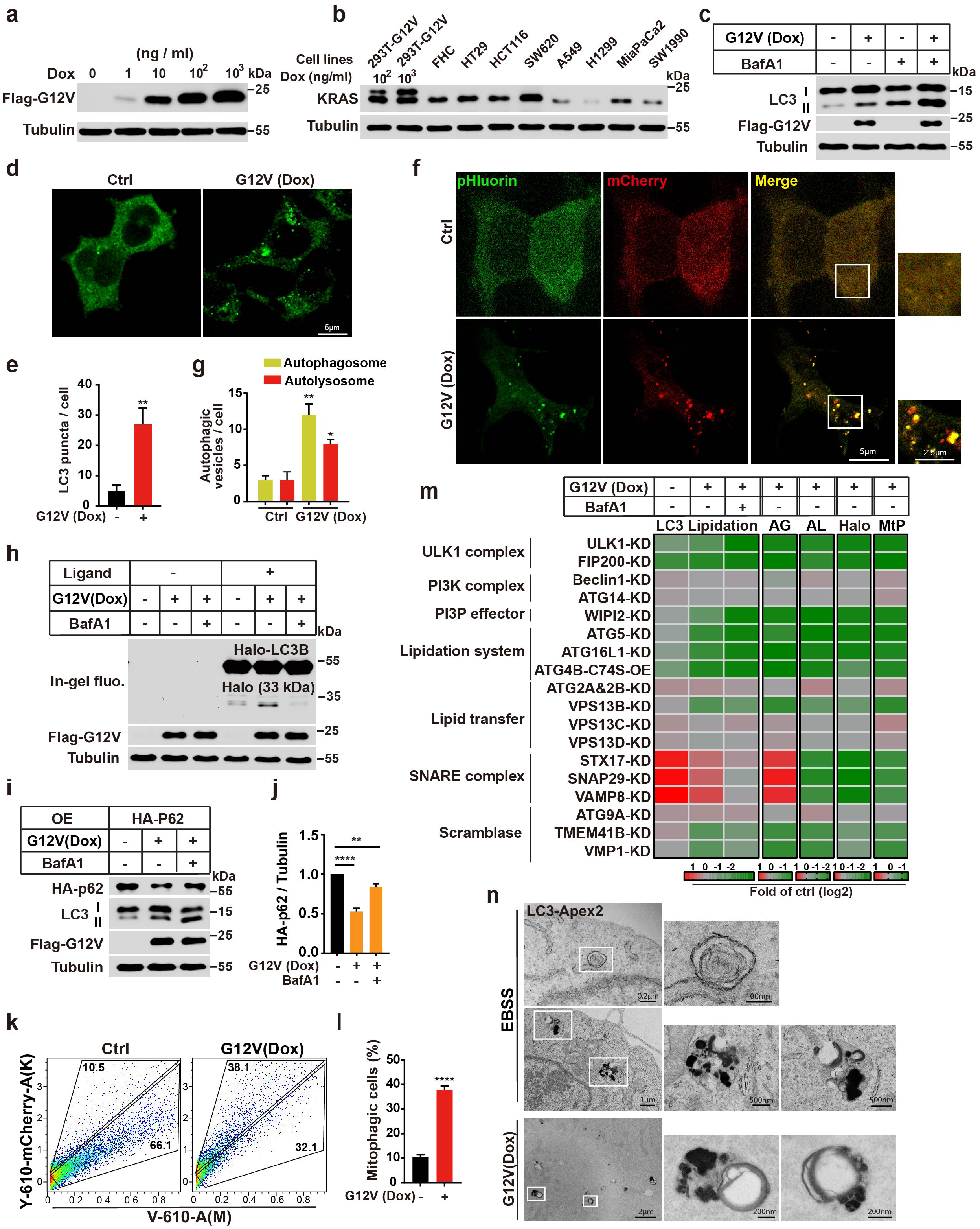
The KRAS(G12V)-induced autophagy system and independency of multiple autophagic factors. **a**. Immunoblot analysis of cell lysates derived from KRAS(G12V) 293T cells treated with a range of Dox concentrations or left untreated for 36 h. **b**. Immunoblot analysis of the expression level of KRAS in different cell lines as indicated. **c**. Immunoblot analysis of LC3 lipidation in control and KRAS(G12V) cells with or without 500 nM Bafilomycin A1 for 1.5 h. **d**.Immunofluorescence and confocal microscopy imaging were performed to visualize the KRAS(G12V)-induced LC3 puncta in 293T cells. Representative cell images are shown. Scale bar sizes are indicated in the image. **e**. Quantification of the LC3 puncta in control and KRAS(G12V) cells in (d) (mean ± SEM). Three experiments (50 cells for each group/experiment) were performed for the statistics (two-tailed t-test). **, *p<*0.01. **f**. Detection of KRAS(G12V)-induced autophagy by the tandem fluorescent LC3 system in control and KRAS(G12V) cells stably expressing mCherry-pHluorin-LC3B. Representative cell images are shown. Scale bar sizes are indicated in the image. **g**. Quantification of the yellow (RFP^+^GFP^+^) and Red (RFP^+^GFP^-^) LC3 puncta. Data are represented as mean ±SEM. Three experiments (50 cells for each group/experiment) were performed for the statistics (two-tailed t-test). *, *p<*0.05; **, *p<*0.01. **h**. Immunoblotting and in-gel fluorescence detection in control and KRAS(G12V) cells stably expressing HaloTag (Halo)-LC3B pulse-labeled for 20 min with 100 nM tetramethyl rhodamine (TMR)-conjugated ligand in nutrient-rich medium with or without 500 nM Bafilomycin A1 for 2 h. **i**. Immunoblot showing the degradation of exogenous p62 in control and KRAS(G12V) cells with or without 500 nM Bafilomycin A1 for 1.5 h. **j**. Quantification of the ratio of P62 to tubulin with the control set as 1.00 (control cells without Bafilomycin A1) analyzed in (k) (mean ± SEM). Three experiments were performed for the statistics (two-tailed t-test). **, *p<*0.01; ****, *p<*0.0001. **k**. FACS analysis of control and KRAS(G12V) cells co-expressing mt-Keima and Parkin using V610 and Y610-mCherry detectors (Beckman CytoFLEX LX). The FACS results are representative of at least three independent experiments. **l**. The percentage of cells with mitophagy based on Y610-mCherry/V610 calculated for (h). Data are represented as mean ±SEM. Three experiments were performed for the statistics (two-tailed t-test). ****, *p<*0.0001. **m**. Heatmap to show the changes of LC3 lipidation (the ratio of lipidated LC3 to tubulin with the control set as 1.00, related to Supplementary information, Fig. S1d, e), autophagic flux by the tandem fluorescent LC3 system (AG (autophagosome) and AL(autolysosome), the control set as 1.00, related to Supplementary information, Fig. S1g, h) and HaloTag-LC3B processing assay (Halo, Halo-TMR band intensity was normalized by the sum of the band intensities Halo-TMR-LC3B and Halo-TMR, and the control set (KRAS(G12V) 293T cells with ligand without Bafilomycin A1) as 1.00. Related to Supplementary information, Fig. S1i, g), and mitophagy (MtP, the control set as 1.00, related to Supplementary information, Fig. S1k, l) in control and KRAS(G12V) 293T cells with knocking down proteins as indicated or overexpressing ATG4B-C74S respectively. Color represents the log2 (fold) of counts per samples. **n**. Electron microscopy images of APEX2-labeled LC3 and the autophagosomes in EBSS or KRAS(G12V) 293T cells. Scale bar sizes are indicated in the image.

autophagosome maturation, indicating that KRAS(G12V) expression activates autophagy (Fig. 1c). Similar results were obtained in 293T cells expressing KRAS(G12D) or KRAS(G12C) alleles (Supplementary information, Fig. S1a). Consistently, LC3 puncta, an indicator of autophagosomes, was increased in Dox-treated KRAS(G12V) cells compared to controls (Fig. 1d, e). In addition, an LC3 paralogue GABARAP colocalized with LC3 in the puncta indicating a general ATG8ylation events on the autophagic membrane (Supplementary information, Fig. S1f).

A double-fluorescence (mCherry-pHluorin) LC3 reporter ^36^ was utilized to confirm that KRAS(G12V) enhanced autophagic flux. In this system, autophagosomes (yellow) and autolysosomes (red) were analyzed by fluorescence. Consistent with the effect on autophagosome biogenesis, Dox induction of KRAS(G12V) increased the yellow and red puncta (Fig. 1f, g), indicating that KRAS(G12V) increased both autophagosome formation and autophagic flux. Furthermore, we employed a HaloTag (Halo)-based reporter processing assay to validate the augmented autophagic flux induced by KRAS(G12V) (Fig. 1h) .^37^ Subsequently, to corroborate autophagic degradation, we assessed P62 clearance and mitophagy using a mito-keima assay (Fig. 1i-l).^38^ Once more, these findings consistently illustrated an enhanced autophagic turnover following KRAS(G12V) expression.

Several inhibitors of KRAS(G12C) have entered clinical trials, including AMG510 (Amgen), which is being evaluated for the treatment of non-small cell lung cancer, colorectal cancer (CRC), and other solid tumors harboring this mutation.^15^ To determine if autophagy induced by mutant KRAS depends on KRAS activation in our system, we tested the effect of AMG510 on LC3-II. The treatment with AMG510 reduced the levels of LC3- II in cells expressing KRAS(G12C), but not in cells expressing KRAS(G12V) or KRAS(G12D) (Supplementary information, Fig. S1b), suggesting that KRAS activation triggers autophagy upregulation.

Two recent work found that RAS activation may play a negative role on autophagy in multiple KRAS-mutant cancer cells but instead MEK inhibition enhanced autophagy in these cells.^39, 40^ To clarify the effect of RAS activation on autophagy, we knocked down RAS across six cancer cell lines with Ras mutation and found that it inhibited LC3 lipidation and mitophagy (Supplementary information, Fig. S2a-c), which supports the positive role of RAS activation on autophagy reported by several other studies,^19, 20, 33–35, 41^. Furthermore, trametinib treatment did not notably upregulate autophagy (Supplementary information, Fig. S2a-c).

### Common and distinct utilization of autophagic factors in KRAS(G12V)- and starvation- induced autophagy

To elucidate the role of autophagic factors in oncogenic KRAS-induced autophagy, we utilized gene knockdown to target key autophagic factors involved in various regulatory steps of starvation-induced autophagy in 293T cells expressing the Dox-inducible KRAS(G12V) construct. These factors encompassed the ULK kinase complex (ULK1 and FIP200), the PI3K complexes (ATG14, and Beclin-1), the PI3P effector WIPI2, the lipidation machinery (ATG5, ATG16L1, and ATG4B-C74S overexpression), the lipid transfer factors ATG2A and ATG2B, ATG9 vesicles (ATG9A), and the SNARE complexes (STX17, SNAP29, and VAMP8). The knockdown efficiency of each gene was validated by western blot or QPCR (Supplementary information, Fig. S1c, d). Autophagic assays were subsequently employed to comprehensively assess the involvement of these factors in KRAS-induced autophagy.

Consistent with the blockade observed in starvation-induced autophagy, inhibition of the ULK kinase complex, the PI3P effector WIPI2, and the lipidation machinery significantly impaired KRAS(G12V)-induced LC3 lipidation (Fig. 1m and Supplementary information, Fig. S1d, e), and autophagic flux as determined by multiple assays (Fig. 1m and Supplementary information, Fig. S1g-l). Moreover, inhibition of the SNARE complex similarly hindered autophagic flux, aligning with observations in starvation-induced autophagy (Fig. 1m and Supplementary information, Fig. S1d, e, g-l). Interestingly, while the PI3K complex, ATG9A, and ATG2s were indispensable for starvation-induced autophagy,^42–51^ their depletion did not impact KRAS-induced autophagy (Fig. 1m and Supplementary information, Fig. S1m-q). Notably, a lipid transfer protein VPS13B ^52, 53^, which functionally transfer lipids akin to ATG2, was essential for KRAS-induced autophagy (Fig. 1m and Supplementary information, Fig. S1d, e, g-l). In addition, two scramblases TMEM41B and VMP1 were involved in KRAS-induced autophagy similar to their role in starvation-induced autophagy (Fig. 1m and Supplementary information, Fig. S1d, e, g-l).^54–59^ These findings collectively suggest RAS-induces a non-canonical autophagy in which it shares and employ distinct sets of autophagic factors (as well as non-autophagic proteins) compared to starvation-induced autophagy. We term it as RAS-induced non-canonical autophagy via ATG8ylation (RINCAA).

The defined set of autophagic factors orchestrates the generation of double-membrane autophagosomes, a hallmark of starvation-induced autophagy. We investigated whether similar double-membrane autophagosomes could be generated with altered autophagic factors. An APEX2 labeling assay, combined with electron microscopy, was performed to determine the structure of APEX2-LC3 labeled membranes. Interestingly, instead of double-membrane autophagosomes generated in starvation-induced autophagy (Fig. 1n), multivesicular and multilaminar structures of LC3 were identified in KRAS(G12V)- expressing cells and multiple cancer cell lines with RAS mutations (Fig. 1n, Supplementary information, Fig. S1r). Notably, irregular multivesicular or multilaminar (occasionally, data not shown) structures of LC3 was also observed in starvation conditions, we consider them as late stages structures such as amphisomes or autolysosomes (Fig.1n). On the contrary RAS-induced multivesicular and multilaminar structure is likely an early stage structure equal to the double-membrane autophagosome (see below Fig. 3e and Supplementary Movie 1). Thus, activated KRAS mutations induce the generation of alternative autophagosomes using a different set of autophagic factors. We term these alternative autophagosomes RAS-induced multivesicular and multilaminar bodies positive of ATG8ylation (RIMMBA).

### PI4KB regulates RINCAA

The intriguing observation of PI3K independence in RINCAA piqued our interest, as the PI3K complex is traditionally considered pivotal in linking protein signaling to membrane remodeling processes in starvation-induced autophagy. Despite the dispensability of PI3K, we found that the PI3P effector WIPI2 remains indispensable in this context (Fig. 1m and Supplementary information, Fig. S1d, e, g-l). Given WIPI2’s potential ability to bind various types of PI-phosphates, we hypothesized that another PI-kinase might assume the role of PI3K in RINCAA KRAS-induced atg8ylation. To identify the specific PI-kinase(s) and PI-phosphates involved, we assessed the impact of different phosphokinase inhibitors on LC3-II levels in KRAS(G12V)-induced 293T cells.

Remarkably, treatment with PI3K inhibitors SAR405,^60^ PIK-III,^61^ VPS34-IN1,^62, 63^ and wortmannin^64^ at 20 nM did not reduce LC3-II levels in Dox-induced cells compared to the control, nor did the PI5K inhibitor YM201636^65^ (Fig. 2a and Supplementary information, Fig. S3a,b). Conversely, these inhibitors effectively attenuated starvation-induced and glucose starvation-induced autophagy, serving as positive controls (Fig. 2a and Supplementary information, Fig. S3c-e). Intriguingly, treatment with 200 nM wortmannin, which inhibits type III PI4Ks^66^, led to decreased LC3-II levels during Dox treatment (Fig. 2a and Supplementary information, Fig. S3b-f), suggesting potential involvement of PI4K and PI4P in RINCAA.

**Fig. 2.**
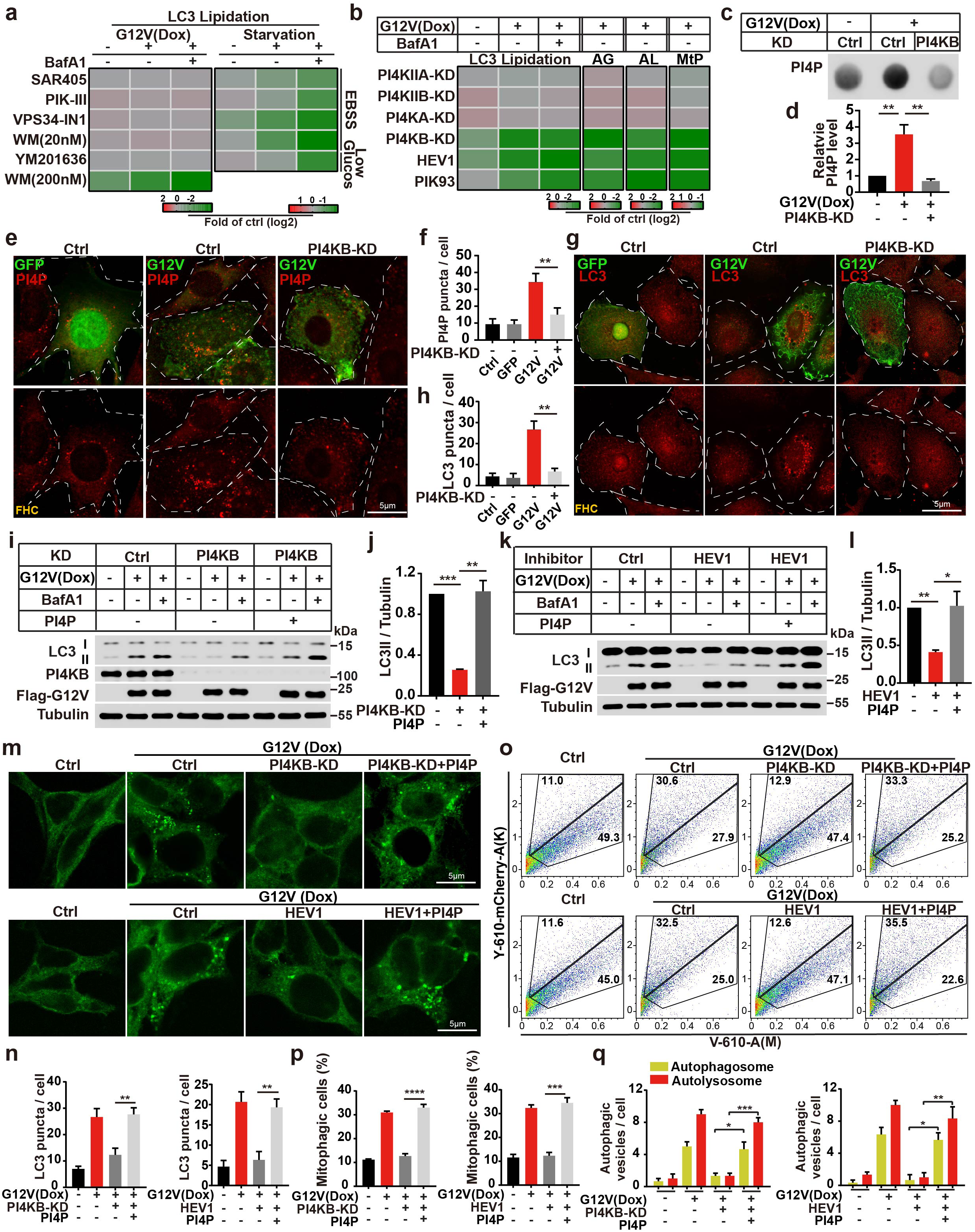
PI4KB regulates RINCAA. **a**. Heatmap to show the changes of LC3 lipidation (the ratio of lipidated LC3 to tubulin with the control set as 1.00) in KRAS(G12V) and starvation groups with different inhibitors (related to Supplementary information, Fig. S3a-e). Color represents the log2 (fold) of counts per samples. **b**. Heatmap to show the changes of LC3 lipidation (the ratio of lipidated LC3 to tubulin with the control set as 1.00, Supplementary information, Fig. S3h, i), autophagic flux (AG and AL, related Supplementary information, Fig. S3g, k) and mitophagy (MtP, related to Supplementary information, Fig. S3l, m) in KRAS(G12V) 293T cells with knocking down PI4Ks or treatment of PI4KB inhibitors. Color represents the log2 (fold) of counts per samples. **c**. Dot blot analysis of the KRAS(G12V)-induced PI4P generation by knocking down PI4KB in control and KRAS(G12V) 293T cells. **d**. Quantification of results shown in (c) (mean ±SEM), and the control set as 1.00. Three experiments were performed for the statistics (two-tailed t-test). **, *p<*0.01. **d**. Immunofluorescence analysis of the PI4P puncta in KRAS(G12V) FHC cells with or without knocking down PI4KB. Representative cell images are shown. Scale bar sizes are indicated in the image. **e**. Quantification of the results shown in (e) (mean ±SEM). Three experiments (50 cells for each group/experiment) were performed for the statistics (two-tailed t-test). **, *p<*0.01. **f**. Immunofluorescence and confocal microscopy imaging were performed to visualize the KRAS(G12V)-induced LC3 puncta in FHC cells with or without knocking down PI4KB. Representative cell images are shown. Scale bar sizes are indicated in the image. **g**. Quantification of the LC3 puncta in control and KRAS(G12V) FHC cells in (g). Data are represented as mean ±SEM. Three experiments (50 cells for each group/experiment) were performed for the statistics (two-tailed t-test). **, *p<*0.01. **i-l.** Immunoblot analysis of the LC3 lipidation rescued by PI4P in KRAS(G12V) 293T cells with suppressed PI4KB by knocking down (i, j) or treatment of the inhibitor (k, l). Quantification of the results in (i, k) are shown in (j, l) (mean ± SEM). Three experiments were performed for the statistics (two-tailed t-test). *, *p<*0.05; **, *p<*0.01;***, *p<*0.001. **m**. Immunofluorescence analysis of the LC3 puncta rescued by PI4P in KRAS(G12V) 293T cells with suppressed PI4KB by knocking down or treatment of the inhibitor. Representative cell images are shown. Scale bar sizes are indicated in the image. **n**. Quantification of the results shown in (m) (mean ±SEM). Three experiments (50 cells for each group/experiment) were performed for the statistics (two-tailed t-test). **, *p<*0.01. **o**. FACS analysis of mitophagy rescued by PI4P in KRAS(G12V) 293T cells with suppressed PI4KB by knocking down or treatment of the inhibitor. The FACS results are representative of at least three independent experiments. **p**. The percentage of cells with mitophagy based on Y610-mCherry/V610 calculated for (o). Error bars represent standard deviations of three experiments (mean ±SEM). Three experiments were performed for the statistics (two-tailed t-test). ***, *p<*0.001; ****, *p<*0.0001. **q**. Quantification of the yellow (RFP^+^GFP^+^) and Red (RFP^+^GFP^-^) LC3 puncta (related to Supplementary information, Fig. S3m). KRAS(G12V) 293T cells were rescued by PI4P with suppressed PI4KB by knocking down or treatment of the inhibitor. Data are represented as mean ±SEM. Three experiments (50 cells for each group/experiment) were performed for the statistics (two-tailed t-test). *, *p<*0.05; **, *p<*0.01; ***, *p<*0.001.

**Fig. 3.**
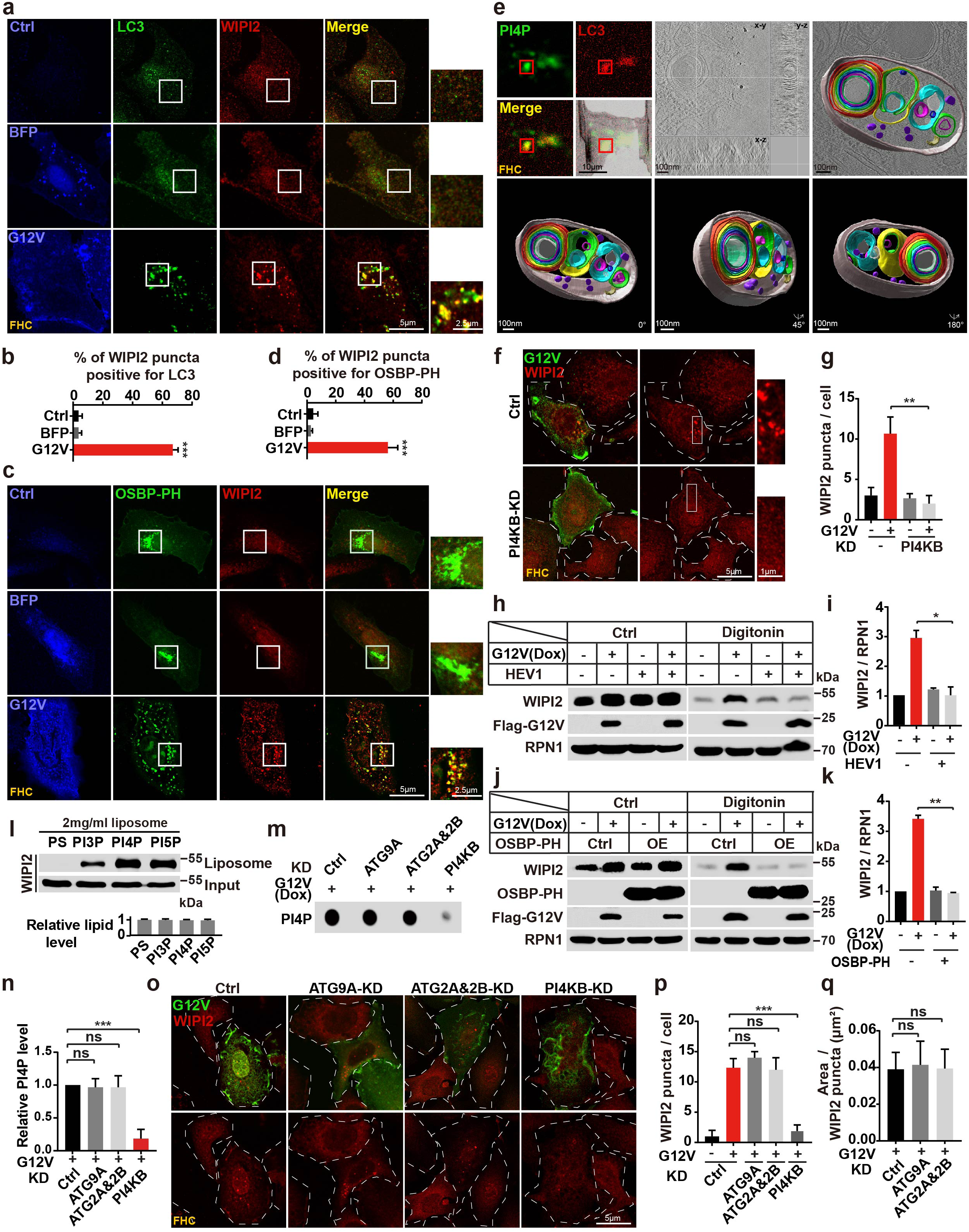
WIPI2 acts as a PI4P effector during RINCAA. **a**. Immunofluorescence and confocal microscopy imaging were performed to visualize the colocalization between the WIPI2 puncta and the LC3 puncta in control and KRAS(G12V) cells. Representative cell images are shown. Scale bar sizes are indicated in the image. **b**. Quantification of the percentage of the WIPI2 puncta colocalizing with the LC3 puncta in (a) (mean ± SEM). Three experiments (50 cells for each group/experiment) were performed for the statistics (two-tailed t-test). ***, *p<*0.001. **c**. Immunofluorescence and confocal microscopy imaging were performed to visualize the colocalization between the WIPI2 puncta and the OSBP-PH puncta in control and KRAS(G12V) cells. Representative cell images are shown. Scale bar sizes are indicated in the image. **d**. Quantification of the percentage of the WIPI2 puncta colocalizing with the OSBP-PH puncta in (c) (mean ±SEM). Three experiments (50 cells for each group/experiment) were performed for the statistics (two-tailed t-test). ***, *p<*0.001. **e**. Light and electron microscopy combined with tomography images of the structure of LC3- and PI4P-positive compartments in KRAS(G12V) FHC cells. Scale bar sizes are indicated in the image. **f**. Immunofluorescence analysis of the KRAS(G12V)-induced WIPI2 puncta with or without knocking down PI4KB. Representative cell images are shown. Scale bar sizes are indicated in the image. **g**. Quantification of the number of the WIPI2 puncta in (f) (mean ± SEM). Three experiments (50 cells for each group/experiment) were performed for the statistics (two-tailed t-test). **, *p<*0.01. **h**. Immunoblot analysis of membrane-bound WIPI2 in control and KRAS(G12V) cells treated with the PI4KB inhibitor T-00127-HEV1 (10 μM). **i**. Quantification of the levels of membrane-bound WIPI2 in (h) (mean ± SEM). Three experiments were performed for the statistics (two-tailed t-test). **, *p<*0.01. **j**. Immunoblot analysis of membrane-bound WIPI2 in control and KRAS(G12V) cells overexpressing OSBP-PH. **k**. Quantification of the levels of membrane-bound WIPI2 in (j) (mean ± SEM). Three experiments were performed for the statistics (two-tailed t-test). **, *p<*0.01. **l**. The liposome flotation assay showing the binding of WIPI2 to PI3P, PI4P, and PI5P. The total lipid level of liposomes was normalized by PC. **m**.Dot blot analysis of the KRAS(G12V)-induced PI4P generation in KRAS(G12V) 293T cells with or without knocking down ATG9A, ATG2A&2B or PI4KB. **n**. Quantification of the results in (m) (mean ±SEM). Three experiments were performed for the statistics (two-tailed t-test). ***, *p<*0.001. **o**. Immunofluorescence analysis of the KRAS(G12V)-induced WIPI2 puncta with knocking down ATG9A, ATG2A&2B or PI4KB in FHC cells. Representative cell images are shown. Scale bar sizes are indicated in the image. **p**. Quantification of the number of the WIPI2 puncta in (o) (mean ± SEM). Three experiments (50 cells for each group/experiment) were performed for the statistics (two-tailed t-test). ***, *p<*0.001. **q**. Quantification of areas of the WIPI2 puncta in (o) (mean ±SEM). Three experiments (50 cells for each group/experiment) were performed for the statistics (two-tailed t-test).

We evaluated the LC3-II levels and autophagic flux assays in our system with shRNA-mediated knockdown of PI4K isoforms to identify which PI4K is the main regulator of RINCAA. The knockdown efficiency of all isoforms was > 80% (Supplementary information, Fig. S3c). Knockdown of PI4KIIA, PI4KIIB, and PI4KA did not affect KRAS(G12V)-induced LC3-II accumulation, whereas PI4KB knockdown resulted in marked depletion of LC3-II and autophagic flux (Fig. 2b and Supplementary information, Fig. S3h-m). We examined the effect of the PI4KB inhibitors on RINCAA to confirm the effects of PI4KB. As a result, T-00127-HEV1^67^ and PIK93^68, 69^ robustly inhibited KRAS(G12V)-induced LC3 lipidation and autophagic flux (Fig. 2b and Supplementary information, Fig. S3h-m). Furthermore, we assessed the PI4P levels using dot blot and PI4P staining assays and found that KRAS(G12V) expression increased PI4P levels which were abrogated by PI4KB knockdown (Fig. 2c-f). Similarly, PI4KB silencing or the T-00127- HEV1 treatment markedly reduced KRAS(G12V)-induced LC3 lipidation and puncta (both FHC expressing the KRAS(G12V) and Dox-induced KRAS(G12V) HEK293T), as well as autophagic turnover in the Dox-induced KRAS(G12V) HEK293T (Fig. 2g-q). And adding exogenous PI4P reversed this effect in the Dox-induced KRAS(G12V) HEK293T (Fig. 2i-q). The data indicate that oncogenic KRAS induces RINCAA through a PI4P- dependent mechanism and PI4KB-mediated generation of PI4P can substitute for the canonical PI3K function during RINCAA.

### Inhibiting PI4KB decreases the proliferation of tumor cells with RAS mutations

KRAS-induced autophagic flux supports the survival of tumor cells.^70–73^ To determine the role of PI4KB in regulating RINCAA and cell proliferation in tumors, we treated two CRC cell lines with different KRAS mutations, such as HCT116 (KRAS^G13D^) and SW620 (KRAS^G12V^), with PIK93 or in combination with the MEK inhibitor trametinib. Consistent with the previous work,^74^ trametinib reduced the proliferation of HCT116 and SW620 cells *in vitro*. Additionally, we observed inhibited cell proliferation by PIK93 and a combined effect with trametinib (Supplementary information, Fig. S4a, b). We next used a subcutaneous xenograft tumor model and the two KRAS mutated cell lines in NOD/SCID mice. Tumors were treated with the control, trametinib, PIK93, or a combination. The animals were followed to monitor tumor size. Consistent with the cellular data, trametinib and PIK93 treatment alone slowed tumor growth in KRAS-mutant tumors, and the drug combination further inhibited tumor growth (Supplementary information, Fig. S4c-h). Immunofluorescence and immunohistochemistry analyses revealed decreased LC3 puncta and increased P62 in PIK93-treated mice and decreased p-ERK in tumors from trametinib-treated mice. The cell proliferation marker Ki67 was decreased in the single-treated group and further decreased in the double-treated group (Supplementary information, Fig. S4i-l). These data indicate that inhibiting PI4K reduced tumor cell autophagy and proliferation, and this strategy can be combined with inhibiting MEK to enhance the tumor inhibitory effect, which is consistent with several studies showing the beneficial effect of inhibiting autophagy together with inhibiting MEK when treating cancer.^18, 39, 40^

### WIPI2 acts as a PI4P effector in RINCAA

The mammalian orthologue of yeast Atg18, WIPI2, is a WD40-repeat-containing PI3P- binding protein initially reported to facilitate LC3 lipidation and the subsequent growth of the phagophore by recruiting the ATG12–ATG5/ATG16L1 complex. ^75^ In addition to binding PI3P, WIPI2 interacts with PI4P and PI5P.^76, 77^ Because our data indicate that PI4KB, PI4P, and WIPI2 are required for RINCAA, we sought to test whether WIPI2 is the PI4P effector. WIPI2 formed puncta, which colocalized with LC3 upon KRAS(G12V) induction, indicating that WIPI2 localizes on the RIMMBA during RINCAA (Fig. 3a, b). The WIPI2 puncta also overlapped with the PH domain of OSBP (OSBP-PH which binds to PI4P) with an optimal expression level, suggesting a link between PI4P, WIPI2, and autophagic membranes (Fig. 3c, d).

The data identified RIMMBA as alternative autophagosomes in RAS-induced autophagy. Using correlative light and electron microscopy combined with tomography, we characterized the structure of LC3- and PI4P-positive compartments. Electron tomography consistently demonstrated an autophagosome-sized structure with a limiting membrane enclosing multivesicular and multilaminar vesicles (Fig. 3e and Supplementary Movie 1). Therefore, these findings establish a connection between RIMMBA and the presence of PI4P and LC3, suggesting that RIMMBA is an early structure of RINCAA equivalent to the double-membrane autophagosome in canonical autophagy.

We performed the following experiments to determine if the location of WIPI2 on RIMMBA was dependent on PI4P. In one experiment, we knocked down PI4KB, which decreased the number of WIPI2 puncta induced by KRAS(G12V) (Fig. 3f, g). In the other two experiments, we reduced PI4P via T-00127-HEV1 treatment or limited the accessibility of PI4P by overexpressing OSBP-PH. In both conditions, decreasing the level or accessibility of PI4P reduced the amount of membrane-bound WIPI2 as revealed by a membrane isolation assay (Fig. 3h-k). Taken together, these data indicate that PI4P is required for WIPI2 targeting to the autophagic membrane.

To confirm WIPI2 binding, we analyzed the binding capacity between WIPI2 and PI4P using a lipid flotation assay. The results showed that WIPI2 bound to PI4P with a higher capacity than that to PI3P or PI5P (Fig. 3l). In line with the above data of RINCAA analysis, ATG9A and ATG2s were not required for PI4P production and WIPI2 puncta formation with KRAS(G12V) (Fig. 3m-q). ATG9A and ATG2s were not involved in regulating PI4P levels in multiple cancer cell lines with RAS mutations (Supplementary information, Fig. S6O). Collectively, these data indicate that WIPI2 acts as a PI4P effector in RINCAA.

### The P38**-**ULK1**-**PI4KB**-**WIPI2 cascade regulates RINCAA

RAS activates the ERK, PI3K/AKT, P38, JNK, and Ral pathways, resulting in distinct cellular responses (Fig. 4a).^10, 78–80^ To examine which of these downstream pathways is involved in RINCAA, we tested a series of specific inhibitors. The effect of the inhibitors was validated by their targets (Supplementary information, Fig. S5a). The P38-specific inhibitor SB203580 ^81^ reduced LC3 lipidation in Dox-induced cells compared to the control, whereas inhibitors of ERK (FR180204)^82^, PI3K/AKT (MK2206)^83^, and Ral (RBC8)^84^ did not (Fig. 4b, Supplementary information, Fig. S5b,c). The effect of inhibiting P38 on RINCAA was further confirmed by the autophagic turnover assays including the double-fluorescence LC3 assay and the mito-keima assay (Fig. 4b, Supplementary information, Fig. S5d-g). The JNK inhibitor JNK-IN-8^85^ increased LC3 lipidation upon KRAS(G12V) expression but reduced LC3 lipidation after treatment with Bafilomycin A1, suggesting an effect on the maturation of the RIMMBA. This notion was confirmed by autophagic turnover assays, in which inhibiting JNK did not affect the yellow puncta but decreased the red puncta in the double-fluorescence LC3 assay (Supplementary information, Fig. S5d-e). JNK inhibition also decreased mitophagy in the mito-keima assay (Supplementary information, Fig. S5f-g).

**Fig. 4.**
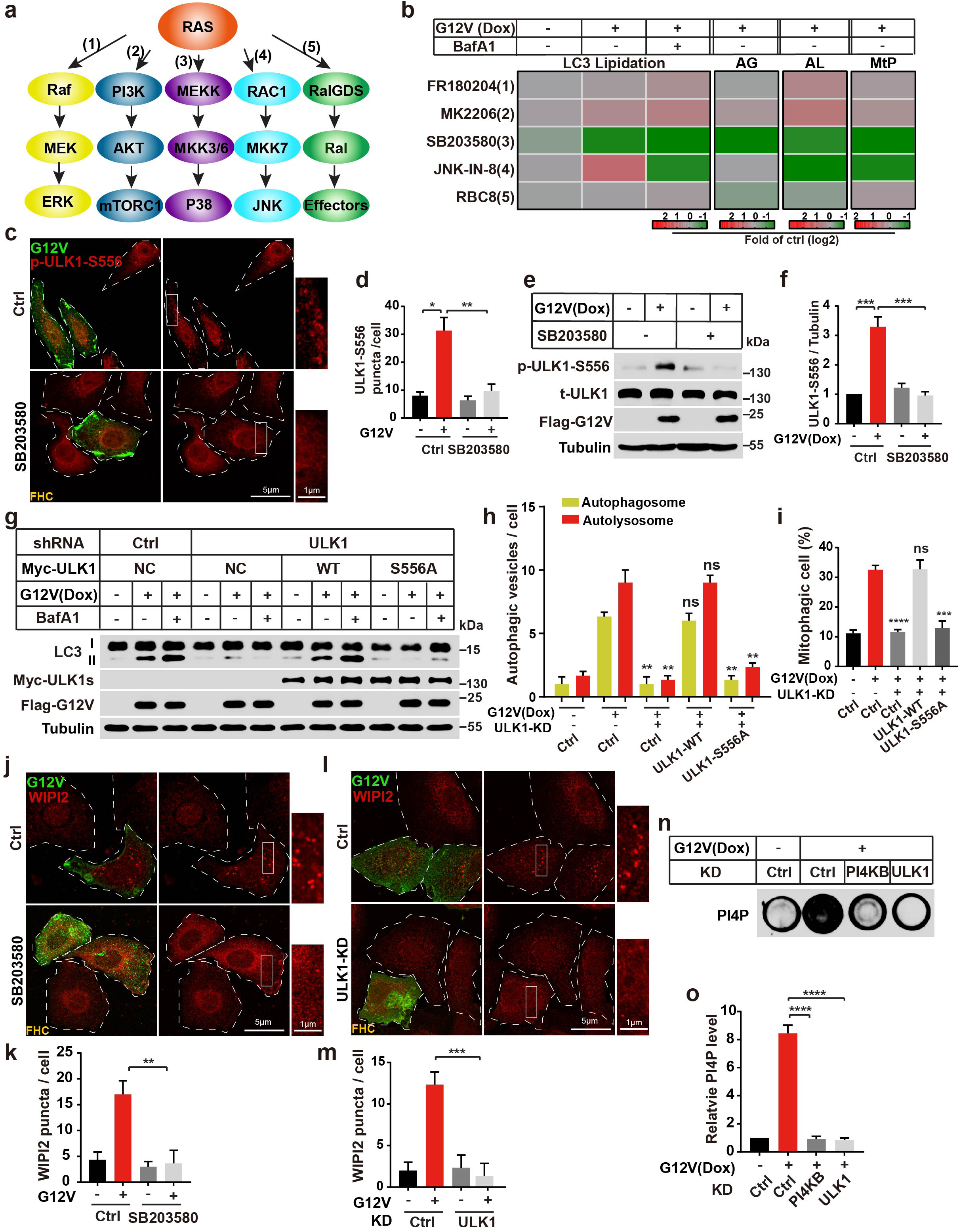
The P38-ULK1-PI4K-WIPI2 cascade regulates RINCAA. **a**. The schematic diagram of the RAS signaling pathway. **b**. Heatmap to show the changes of LC3 lipidation (related to Extended Data Fig. S5b, c), autophagic flux (AG and AL, related to Supplementary information, Fig. S5d, e), and mitophagy (MtP, related to Supplementary information, Fig. S5f, g) in control and KRAS(G12V) 293T cells with different inhibitors in the RAS signaling pathway. Color represents the log2 (fold) of counts per samples. **c**. Immunofluorescence analysis of the regulation of SB203580 on the KRAS(G12V)- induced p-ULK1-S556 puncta. Representative cell images are shown. Scale bar sizes are indicated in the image. **d**. Quantification of the numbers of WIPI2 puncta in (c) (mean ± SEM). Three experiments (50 cells for each group/experiment) were performed for the statistics (two-tailed t-test). *, *p<*0.05; **, *p<*0.01. **e**. Immunoblot analysis of the regulation of SB203580 on the KRAS(G12V)-induced p- ULK1-S556 level. **f**. Quantification of the results in (e) (mean ±SEM). Three experiments were performed for the statistics (two-tailed t-test). ***, *p<*0.001. **g**. Immunoblot analysis of the rescue effects of WT-ULK1 and ULK1-S556A on the KRAS(G12V)-induced LC3 lipidation when knocking down ULK1. **h**. Quantification of the yellow (RFP^+^GFP^+^) and Red (RFP^+^GFP^-^) LC3 puncta (related to Supplementary information, Fig. S5h) to analyze the rescue effects of WT-ULK1 and ULK1-S556A on the KRAS(G12V)-induced LC3 flux in 293T cells. Data are represented as mean ±SEM. Three experiments (50 cells for each group/experiment) were performed for the statistics (two-tailed t-test). **, *p<*0.01. **i**. Quantification of results shown in (Supplementary information, Fig. S5i). The percentage of cells with mitophagy based on Y610-mCherry/V610. Data are represented as mean ±SEM. Three experiments were performed for the statistics (two- tailed t-test). ***, *p<*0.001; ****, *p<*0.0001. **j**. Immunofluorescence analysis of the regulation of SB203580 on the KRAS(G12V)- induced WIPI2 puncta in FHC cells. Representative cell images are shown. Scale bar sizes are indicated in the image. **k**. Quantification of the numbers of WIPI2 puncta in (j) (mean ± SEM). Three experiments (50 cells for each group/experiment) were performed for the statistics (two-tailed t-test). **, *p<*0.01. **l**. Immunofluorescence analysis of the KRAS(G12V)-induced WIPI2 puncta with knocking down ULK1 in FHC cells. Representative cell images are shown. Scale bar sizes are indicated in the image. **m**. Quantification of the numbers of WIPI2 puncta in (l) (mean ± SEM). Three experiments (50 cells for each group/experiment) were performed for the statistics (two-tailed t-test). ***, *p<*0.001. **n**. Dot blot analysis of the KRAS(G12V)-induced PI4P generation with knocking down PI4KB or ULK1 in 293T cells. **o**. Quantification result for (n) (mean ±SEM). Three experiments were performed for the statistics (two-tailed t-test). ****, *p<*0.0001.

Because inhibiting P38 blocks the biogenesis of RIMMBA, we next focused on this pathway. The P38-MAPK pathway mediates the activation of autophagy by activating mouse ULK1 through phosphorylation on serine (S) 555.^86^ We observed the formation of ULK1-S556 puncta (an indicator of ULK1 phosphorylation and early autophagosome formation, human ULK1 S556 corresponds to mouse ULK1 S555) upon KRAS(G12V) expression, which was mitigated by inhibiting P38 (Fig. 4c, d). Immunoblot analysis confirmed the increase in ULK-S556 phosphorylation, which was inhibited by SB203580 (Fig. 4e, f). We next performed rescue experiments in ULK1-depleted cells. Restoration of wild-type ULK1 but not of a ULK1-S556A point mutant restored LC3 lipidation upon KRAS(G12V) expression (Fig. 4g-i, Supplementary information, Fig. S5h, i), indicating that P38-mediated activation of ULK1 on S556 is important for RINCAA. The requirement of P38 activation in RINCAA was also confirmed in multiple cancer cell lines (HCT116, SW620, H1299, A549, MiaPaCa2, and SW1990) with endogenous RAS mutation (Supplementary information, Fig. S5j-l). Of notice, another study also highlighted the significance of ULK1 S556 phosphorylation in activating autophagy in RAS mutant cancer cell lines. This study demonstrated that an MEK inhibitor enhances LKB1 and AMPK1 activation, subsequently leading to ULK1 S556 phosphorylation and the enhancement of autophagy.^40^

Next, we determined the relationship between P38, ULK1, PI4P, and WIPI2. SB203580 treatment or ULK1 knockdown inhibited the formation of WIPI2 puncta induced by KRAS(G12V) expression (Fig. 4j-m). ULK1 knockdown blocked the increase in PI4P induced by KRAS(G12V) expression similar to PI4KB depletion (Fig. 4n, o). These data indicate that P38 activates ULK1 via S556 phosphorylation and ULK1 activates PI4P biogenesis and WIPI2 targeting to early RIMMBA structures in the context of oncogenic KRAS signaling.

### ULK1-mediated PI4KB phosphorylation is required for RINCAA and KRAS-mutated tumor growth

To understand how activating ULK1 affects PI4KB, we conducted a Phos-tag gel assay, in which KRAS(G12V) overexpression caused a gel mobility shift in PI4KB to a higher molecular weight, suggesting more phosphorylation (Fig. 5a). Interestingly, ULK1 knockdown reversed the mobility shift of PI4KB upon KRAS(G12V) expression (Fig. 5b). In addition, ULK1 was associated with PI4KB in a co-immunoprecipitation experiment (Fig. 5c). Therefore, these data indicate that ULK1 is involved in PI4KB phosphorylation. Mass spectrometry analysis identified five PI4KB phosphorylated peptides in which phosphorylation of Peptide-1 (K251-K269) indicated regulation by ULK1 activity (Fig. 5d and Supplementary information, Table S1). This peptide contains S256, S258, T263, and S266 as potential ULK1 phosphorylation sites.

**Fig. 5.**
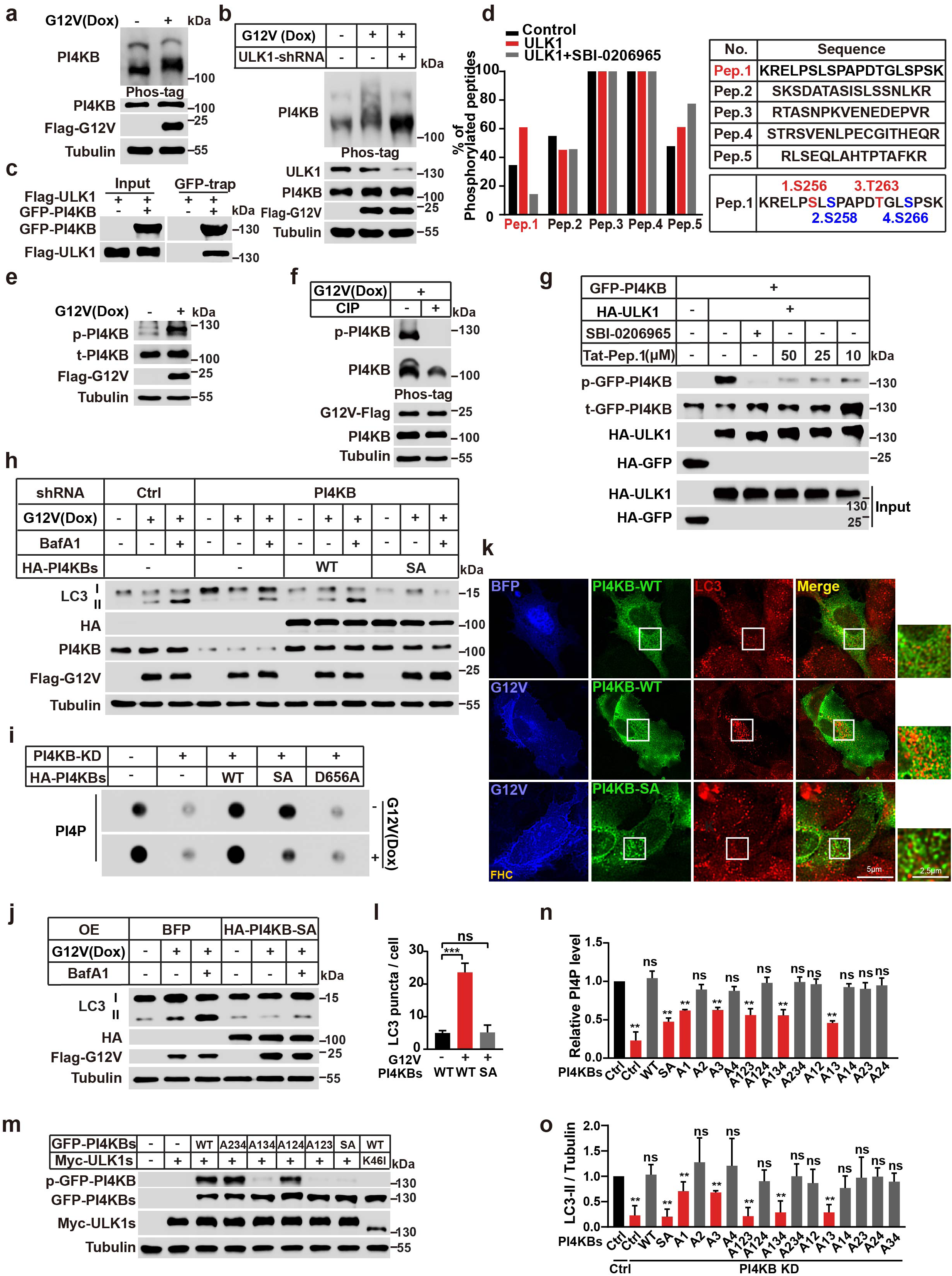
ULK1 mediates PI4KB phosphorylation on the region of K251-K269 which is required for RINCAA. **a**. Immunoblot analysis of the PI4KB phosphorylation from control and KRAS(G12V) 293T cells by Phos-tag gel. **b**. Immunoblot analysis of the PI4KB phosphorylation from control and KRAS(G12V) 293T cells with or without knocking down ULK1 by Phos-tag gel. **c**. Co-immunoprecipitation analysis for the interaction between PI4KB and ULK1. **d**. Mass spectrometry analysis of the percentage of the five phosphorylated PI4KB peptides. **e**. Immunoblot analysis of the PI4KB phosphorylation by the antibody against p-PI4KB in control and KRAS(G12V) 293T cells. **f**. Immunoblot analysis of the PI4KB phosphorylation by the antibody against p-PI4KB in KRAS(G12V) 293T cells treated with or without CIP. **g**. Immunoprecipitation and in vitro kinase assay to analyze the PI4KB phosphorylation by ULK1. HEK293T cells were transfected with HA-tagged ULK1 constructs, lysed, and used for immunoprecipitation reactions. Immunoprecipitates were incubated in an in vitro kinase reaction mixture containing ATP and the substrate GFP-PI4KB with or without SBI-0206965 (10 μM) and Pepptide1(Tat-Pep.1) as indicated concentration. Reaction products were resolved on SDS-PAGE gels. Anti-HA immunoblotting was performed as a control to quantify the amount of ULK1 protein precipitated. The PI4KB phosphorylation was detected by the antibody against p-PI4KB. The experiment was reproduced three times. **h**. Immunoblot analysis of the rescue effect of PI4KB-WT or PI4KB-SA on KRAS(G12V)-induced LC3 lipidation after knocking down PI4KB in 293T cells in the absence or presence of 500 nM Bafilomycin A1 for 1.5 h. **i**. Dot blot analysis of the effect of PI4KB mutants on the KRAS(G12V)-induced PI4P generation in 293T cells. **j**. Immunoblot analysis of the KRAS(G12V)-induced LC3 lipidation when overexpressing PI4KB-SA in 293T cells in the absence or presence of 500 nM Bafilomycin A1 for 1.5 h. **k**. Immunofluorescence analysis of the KRAS(G12V)-induced LC3 puncta when overexpressing PI4KB-SA in FHC cells. Representative cell images are shown. Scale bar sizes are indicated in the image. **l**. Quantification of the LC3 puncta in (k) (mean ±SEM). Three experiments (50 cells for each group/experiment) were performed for the statistics (two-tailed t-test). ***, *p<*0.001. **m**. Immunoblot analysis changes of the p-PI4KB by the different site mutation of PI4KB S256(1), S258(2), T263(3) and S266(4) in control and ULK1 overexpressing 293T cells. **n**. Quantification of PI4P level by the different site mutation of PI4KB-Peptide.1 in KRAS(G12V) 293T cells of (Supplementary information, Fig. S6d) (mean ± SEM). Three experiments were performed for the statistics (two-tailed t-test). **, *p<*0.01. **o**. Statistical analysis of LC3 lipidation by overexpressing different site mutation of PI4KB S256(1), S258(2), T263(3) and S266(4) in control and PI4KB-knockdown KRAS(G12V) 293T cells of (Supplementary information, Fig. S6e) (mean ± SEM). Three experiments were performed for the statistics (two-tailed t-test). **, *p<*0.01.

To further elucidate PI4KB phosphorylation at this region, we developed a phosphorylation-specific antibody targeting its four potential phosphorylation sites. Upon induction of KRAS(G12V) expression, we observed an augmented phosphorylation signal, which was abrogated by treatment with calf-intestinal alkaline phosphatase, confirming the antibody’s specificity (Fig. 5e, f). The antibody’s specificity was further validated in HCT116, SW620 and MiaPaCa2, where the phosphorylation signal in immune-blot or immune-staining was inhibited by RAS knockdown or AMG510, respectively (Supplementary information, Fig. S6a-c). Utilizing this antibody, we investigated whether ULK1 could directly phosphorylate potentially at S256, S258, T263, and S266 within the context of PI4KB. Immune-isolated ULK1 and PI4KB were subjected to an in vitro reaction with ATP. Our results indicate that the phosphorylation occurs in the presence of ULK1, an effect that was fully abrogated by treatment with the ULK1 inhibitor SBI-0206965. Additionally, excessive synthesized Peptide-1 (K251-K269 of PI4KB), likely competing with PI4KB for ULK1 phosphorylation, also abolished the phosphorylation (Fig. 5g).

To determine the function of PI4KB phosphorylation on region K251-K269, we generated a phosphorylation-deficient mutant by mutating the four potential phosphorylation sites to alanine (A) (SA mutant). Re-expression of WT PI4KB in PI4KB knockdown cells restored LC3 lipidation upon KRAS(G12V) expression, whereas the SA mutant did not (Fig. 5h), suggesting that phosphorylation of region K251-K269 is required for PI4KB to regulate RINCAA. To analyze the effect of the SA mutation on kinase activity, we performed a PI4P dot blot assay. Knockdown of PI4KB in control cells decreased PI4P, which was restored by the WT and SA PI4KB but not by the kinase-dead mutant (D656A). However, WT PI4KB restored the upregulated PI4P but the SA mutant failed to enhance PI4P production upon PI4KB depletion in KRAS(G12V) expressing cells (Fig. 5i). Taken together, these data suggest that the phosphorylation of region K251-K269 may be a form of PI4KB kinase activity that specifically regulates RINCAA.

In addition to loss of function, the PI4KB SA mutant generated a dominant-negative effect on RINCAA when overexpressed, as revealed by LC3 lipidation and puncta formation (Fig. 5j-l). Consistently, overexpressing PI4KB-SA inhibited RINCAA and cell proliferation in the RAS mutant cell lines (HCT116, SW620, H1299, A549, MiaPaCa2, and SW1990) (Fig. 6a-c). Therefore, we subsequently determined the anti-neoplastic effects of PI4KB-SA expression using the subcutaneous xenograft tumor model. We generated HCT116 and SW620 cell lines stably overexpressing PI4KB-SA which were employed in the xenograft tumor experiment. Consistent with inhibiting PI4KB, PI4KB- SA expression reduced RINCAA (as revealed by LC3 and P62 staining), cell proliferation (Ki67), and tumor growth. The effect was combined with the MEK inhibitor treatment (Fig. 6d-h). These findings indicate that ULK1-mediated PI4KB phosphorylation is required for RINCAA and tumor growth.

**Fig. 6.**
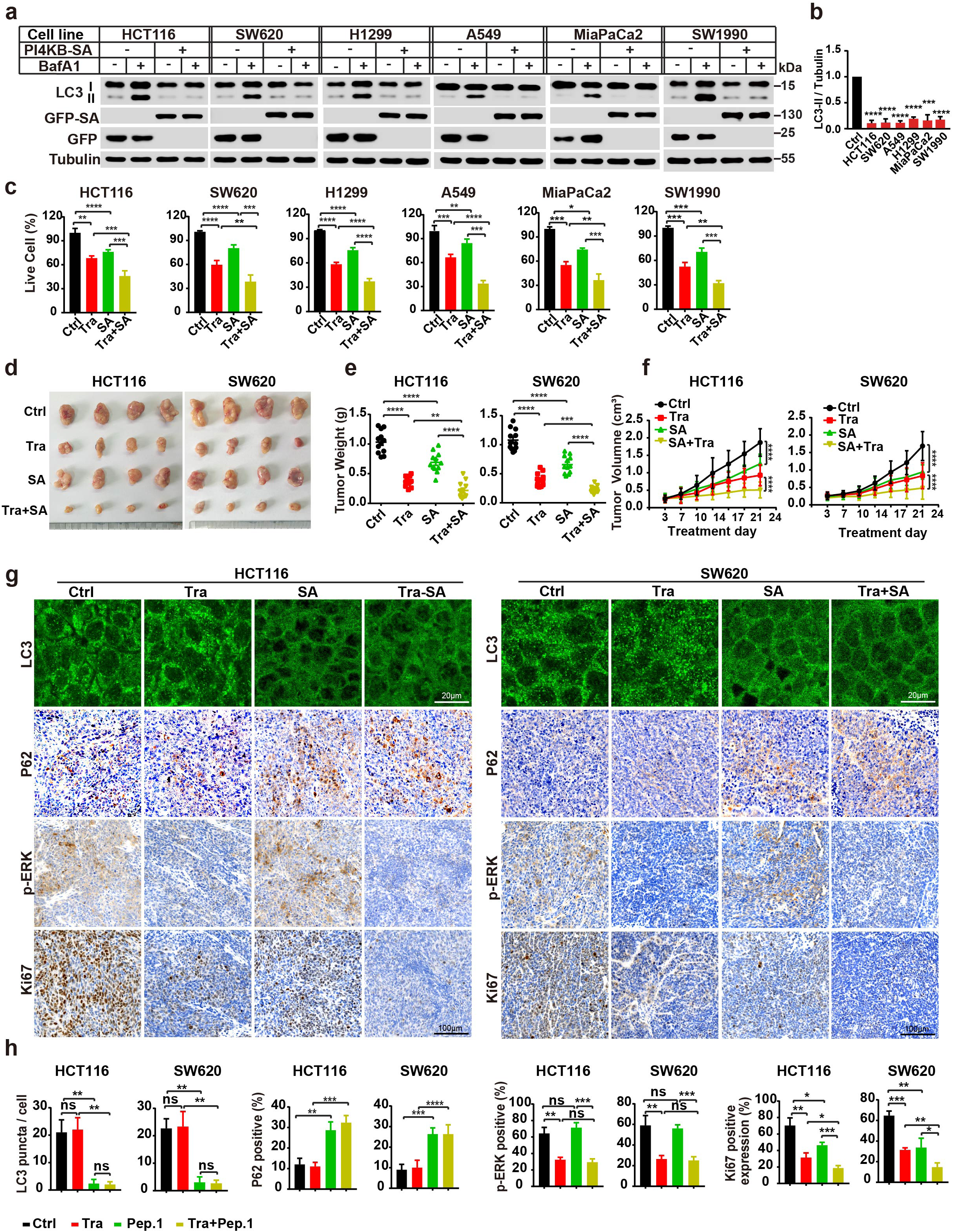
Inhibitory effect of PI4KB-SA on RAS-mutant cell proliferation and xenograft tumor. **a**. Immunoblot analysis of the inhibitory effect of the PI4KB-SA on LC3 lipidation in the control and RAS-mutant cancer cell lines in the absence or presence of 500 nM Bafilomycin A1 for 1.5 h. **b**. Quantification of the ratio of lipidated LC3 to tubulin with the control set as 1.00 (control cells with Bafilomycin A1) analyzed in (d) (mean ±SEM). Three experiments were performed for the statistics (two-tailed t-test). ***, *p<*0.01; ****, *p<*0.0001. **c**. The CCK8 analysis of the cell proliferation of the control and Ras-mutant cell lines treated with the GFP-Peptide 1 or combined with Trametinib (Tra, 100 nM) for 96 h (mean ±SEM). Three experiments were performed for the statistics (two-tailed t-test). **d**. *p<*0.05; **, *p<*0.01; ***, *p<*0.001; ****, *p<*0.0001. **e**. Images of xenograft tumors of HCT116 and SW620 cells from mice treated with: (1) vehicle (Ctrl); (2) Trametinib (Tra); (3) PI4KB-SA overexpression (SA); or (4) the combination of both (Tra+SA) (n = 12 mice (one tumor/mice) in each group). The tumors were removed and photographed after 24-day treatment. **e**. Weights of the xenograft tumors in (g) (means ± SD). The statistical analysis was performed by two-tailed t-test. *, *p<*0.05; **, *p<*0.01; ****, *p<*0.0001. **f**. The growth curve of the xenograft tumors in (g) (means ±SD). Statistical analysis was performed by two-way-ANOVA; ****, *p<*0.0001. **g**. Immunofluorescence and immunohistochemical analysis of the sections of xenograft tumors in (g). Sections were stained with antibody against LC3, P62, p-ERK1/2 or Ki67, as indicated. Scale bar sizes are indicated in the image. **h**. Statistical analysis of the numbers of LC3 puncta, P62 positive rates, p-ERK positive rates, and levels of Ki67 expression in (j) (means ± SD). Statistical analysis was performed by two-tailed t-test; *, *p<*0.05; **, *p<*0.01; ***, *p<*0.001.

The mass spectrometry analysis shown above did not provide precise identification of the phosphorylation sites on Peptide 1. However, through experimentation involving various combinations of phosphorylation site mutations, we discovered that the simultaneous mutation of sites 1 and 3 (S256 and T263) instead of the other two possible sites resulted in the abolition of PI4KB phosphorylation (Fig.5m). Such mutation also impaired the increase of PI4P mediated by PI4KB and compromised RINCAA in the presence of KRAS(G12V) expression (Fig.5 n, o and Supplementary information, Fig. S6d, e). These findings suggest that S256 and T263 are the primary sites of PI4KB phosphorylation crucial for RINCAA activation. A preferred amino acid sequence for ULK1 substrates, characterized by a Leu or Met residue at position −3, along with aliphatic and aromatic hydrophobic residues at positions +1 and +2, has been identified.^87^ The neighboring residues of S256 and T263 contains an L at position +1 and +2 respectively and therefore partially match this sequence.

### ULK1-regulated PI4KB phosphorylation may be a target for treating RAS-mutant cancer

The data indicate that ULK1-regulated PI4KB phosphorylation of S256 and T263 may be a signature of KRAS overactivation and the induction of RINCAA. In immunoblot analysis, cancer cells with a RAS mutation contained a higher level of PI4KB phosphorylation on S256 and T263 than those without a RAS mutation (Fig.7a). In addition, in a mutant-RAS tissue microarray of patients with CRC, the overall PI4KB-Peptide 1 phosphorylation was much higher in RAS-mutated (compared to those with WT RAS) colon or rectal cancer (Fig.7b, c) The data suggest that upregulated PI4KB-Peptide 1 phosphorylation may be a signature of cancer with the RAS mutation.

We investigated the impact of phosphorylation substrate PI4KB-Peptide-1 on ULK1- regulated PI4KB phosphorylation. Consistent with our in vitro ULK1 kinase assay (Fig.5g), expression of PI4KB-Peptide-1 or treating cell with PI4KB-Peptide-1 fused to Tat led to the inhibition of PI4KB phosphorylation on S256 and T263, resulting in decreased PI4P generation and RINCAA blockage as demonstrated by multiple assays, including LC3 lipidation, tandem fluorescence LC3 assay and mitophagy (Fig.7d-l). Therefore, the data confirm the notion that competing against PI4KB phosphorylation on S256 and T263 blocks RINCAA.

We then sought to confirm a similar inhibition of RINCAA by PI4KB phosphorylation on S256 and T263 using cancer cells with endogenous *ras* mutations hindering.

Consistently, PI4KB-Peptide-1-mediated inhibition of PI4KB phosphorylation on S256 and T263 (Fig.8a), and therefore hindered RINCAA, including LC3 lipidation (Fig.8b, c), autophagic flux determined by mitophagy (Fig.8d, e), and WIPI2 puncta formation, across various *ras*-mutated cancer cell lines (Fig.8f, g). Notably, PI4KB-Peptide-1 also suppressed cell proliferation in these cancer cells and exhibited comparable inhibitory effects on proliferation to chloroquine, a clinically used lysosomal inhibitor for autophagy inhibition, when combined with trametinib (Fig.8h).

We employed the tumor xenograft assay mentioned above to determine an inhibitory effect of tumor formation by blocking PI4KB phosphorylation on S256 and T263. In this in vitro tumor model, PI4KB-Peptide-1 expression in combination with trametinib dramatically blocked tumor growth, achieving equivalent (SW620) or superior (HCT116) tumor inhibitory effects to chloroquine combined with trametinib (Fig.9a-c). Immunofluorescence and histochemistry confirmed that PI4KB-Peptide-1 expression curbed PI4KB phosphorylation on S256 and T263, RINCAA, cell proliferation, and tumor growth in the tumor xenograft (Fig.9d, e). Therefore, targeting PI4KB phosphorylation on S256 and T263 specifically inhibits RINCAA in Ras-mutant tumors and suppresses tumor growth.

To further validate the therapeutic potential of inhibiting PI4KB phosphorylation on S256 and T263, we lastly utilized a KPC mice model of pancreatic cancer, known for faithfully recapitulating human pancreatic cancer biology.^88–90^ Consistent with our findings in cancer cell lines and xenograft experiments, treatment with PI4KB Peptide-1 blocked PI4KB phosphorylation on S256 and T263, and RINCAA in cancer tissues and, in combination with trametinib, effectively suppressed tumor growth and prolonged survival (Fig.10a-f). Notably, in contrast to chloroquine, PI4KB Peptide-1 treatment alone decreased tumor growth and increased life span. In addition, the combination of PI4KB Peptide-1 and trametinib exhibited superior efficacy than chloroquine in inhibiting tumor growth, preserving normal pancreatic tissue, and extending survival periods (Fig. 10d-f).

The tumor inhibition effect of PI4KB Peptide-1 surpasses that of chloroquine in immune-competent mice but is similar in immune-deficient mice, suggesting that PI4KB Peptide-1 positively regulates the immune response more effectively than chloroquine.

Recent research demonstrated that autophagy in pancreatic cancer (with ∼98% Ras mutation) impedes immune recognition of cancer cells by degrading MHC-I ^21^. To explore the possible involvement of immune regulation in the observed superior effect of PI4KB Peptide-1, we analyzed MHC-I levels in the tumors (Fig. 10d-f). Both chloroquine and PI4KB Peptide-1 increased MHC-I, consistent with previous findings on autophagy’s role in degrading MHC-I ^21^. However, only PI4KB Peptide-1 enhanced CD8+ T cell infiltration (Fig. 10d-f). The lack of enhanced CD8+ T cell infiltration by chloroquine is likely due to chloroquine’s general inhibition of autophagy in T cells, as autophagy is essential for T cell function ^91–94^. In addition, chloroquine inhibits T cell function ^95, 96^.

Therefore, the tumor-specific inhibition of autophagy by PI4KB Peptide-1 provides a superior tumor-killing effect. This occurs by blocking the metabolic remodeling of cancer cells, enhancing MHC-I-mediated tumor antigen presentation, and minimally inhibiting CD8+ T cell activation. These data further support targeting PI4KB phosphorylation on S256 and T263 as a promising therapeutic strategy against RAS-mutation-driven cancers.

## DISCUSSION

The present study identified a P38-ULK1-PI4KB axis as a special signaling cascade regulating a non-canonical form of autophagy in KRAS mutated cells, which diverges from the nutrient sensing-modulated ULK1-PI3K signaling in starvation-regulated autophagy. Therefore, this study provides implications for how autophagy is specifically regulated under various disease states. Our data suggest a model wherein mutated KRAS activates P38, which in turn triggers the activation of ULK1. ULK1 then phosphorylates PI4KB on S256 and T263 and activates PI4KB to generate PI4P. Finally, WIPI2 acts as a PI4P effector and further recruits the lipidation machinery containing the ATG16L1/ATG12- ATG5 complex to promote the formation of non-canonical autophagic membranes. Differential requirement of PI-kinases, lipid transfer proteins and scramblases etc results in the formation of atypical autophagosomes, termed RIMMBA, instead of the conventional double-membrane autophagosomes (Fig.10).

Although the PI3K complex is an essential player in starvation-induced autophagy by generating PI3P, which converges upstream to the membrane remodeling events,^28, 29, 97, 98^

our study determined that the PI3K complex is not required for RINCAA. Noncanonical PI3K-independent autophagy has been reported elsewhere. Autophagosomes are detected in T lymphocytes and sensory neurons from VPS34^-/-^ mice and in glucose-starved cells treated with the VPS34 inhibitor wortmannin.^99, 100^ PI3K is also not involved in a recently discovered unconventional secretion process regulated by multiple autophagic systems, as well as LC3-dependent EV loading and secretion.^101^ Nonetheless, in several cases, PI- phosphates other than PI3P are generated by specific PI-kinases to regulate the early steps of autophagosome biogenesis. For example, PIKfyve and its product PI5P regulate glucose starvation-induced autophagy^77^ and, in our case, PI4KB catalyzed the formation of PI4P for autophagosome biogenesis in RINCAA. Thus, WIPI2 is a shared effector for PI- phosphates by directly binding and activating the LC3 lipidation process. Therefore, it is likely that the generation of different types of PI-phosphates may represent regulatory diversification employed by different upstream signals converging on WIPI2 to induce autophagosome biogenesis.

PI4P, generated by different PI4Ks, plays multiple roles in different types of autophagy. PI4K2A catalyzes the formation of PI4P on the autophagosome, which facilitates ATG14 recruitment, and autophagosome-lysosome fusion, which is a late step in autophagy.^102–105^ PI4P on the autophagosome is targeted by SteA, which is a *Salmonella* pathogenic factor that blocks the formation of the autolysosome during xenophagy.^106^ PI4P generated by PI4K2A during secretory autophagy regulates the formation of non-canonical autophagosomes, which fuse with the RAB22A-positive early endosome to form a Rafeesome (RAB22A-mediated non-canonical autophagosome fused with an early endosome) that secretes STING via extracellular vesicles.^107^ PI4KA-regulated PI4P production controls autophagosome biogenesis in Arabidopsis.^108^ In addition to PI4K2A, PI4KB generates PI4P, which acts during the early stages of starvation-induced and RINCAA. During starvation, PI4KB is associated with ATG9A and ATG13 during the initial step of autophagy.^109^ Different from the published work, in our study PI4KB replaces the PI3K complex to generate PI4P as a substitute for PI3P in RINCAA, which recruits WIPI2 (Fig.2 and 3).

Our study revealed that ATG2 and ATG9A are dispensable for RINCAA (Fig. 1m). The mechanisms underlying RIMMBA formation in the absence of these proteins. ATG2 functions as a lipid channel facilitating phospholipid delivery to the expanding autophagosome.^45, 49, 50^ Intriguingly, our findings indicate that the lipid channeling protein VPS13B is necessary for RINCAA (Fig. 1m). The involvement of Vps13 in autophagy has been reported in yeast.^52^ Moreover, one member of the VPS13 paralogues VPS13D was shown to regulate mitophagy in mammals.^53^ Therefore, VPS13B may act as substitutes for ATG2 in phagophore expansion. Additionally, ATG9A, a scramblase critical for autophagosome biogenesis, may be functionally replaced by other scramblases such as TMEM41B or VMP1 during RINCAA.^54–59^ Notably, VPS13s, TMEM41B, and VMP1 have been implicated in the regulation of autophagosome biogenesis in previous studies.^52–57^

The coordination between RINCAA and canonical autophagy within the cell remains incompletely understood. Our findings show that treatment with rapamycin and KRAS induction leads to an additive increase in LC3 lipidation, suggesting that these two autophagy pathways can operate concurrently within the cell (Supplementary information, Fig. S6f, g). However, it is plausible that competition may arise between these pathways when shared autophagic components become limited. Notably, under conditions of KRAS induction and starvation, ULK1 demonstrates selective activation of PI4KB and components of the PI3K complex (including Beclin-1), indicating that specific regulatory signals modulate ULK1’s role in coordinating these distinct forms of autophagy (Supplementary information, Fig. S6h).

RAS mutations occur at the highest rate (∼25%) among cancers and are associated with several aggressive cancers.^110^ KRAS mutations are found in approximately 85% of all Ras-driven cancers and are thus regarded as the most frequently mutated oncogene in human cancer.^111–113^ Hyperactive RAS mutants are challenging to target due to the absence of drug-binding pockets on RAS proteins^14^. Until recently, the only clinically effective inhibitors were AMG510 and MRTX849, which specifically target KRAS (G12C).^15,16^ Although several KRAS G12X inhibitors have been developed, comprehensive clinical evaluations are lacking, and patients can develop resistance through feedback mechanisms and genetic alterations in the RAS pathway ^114–116^. Therefore, identifying additional targets in RAS-mutated cancers is essential to improve therapeutic efficacy and address resistance. As an alternative approach, targeting the signaling pathway downstream of RAS has been extensively explored. Autophagy has been reported as a downstream effect of RAS signaling.^19, 117^ High dependence on autophagy has been reported and combined inhibition of autophagy and ERK (activation of another downstream effect of RAS signaling) has been shown by multiple studies as a potential strategy against cancer with a RAS mutation.^18, 39, 40^ Nonetheless, physiological autophagy also plays essential roles in maintaining cellular homeostasis and protecting cells against stress, such as starvation.^3, 118^ Therefore, it is necessary to identify the regulatory differences between physiological and RAS-related autophagy for specific and efficacious targeting of autophagy in cancers with RAS mutations. Our study identified differential regulation in the PI-phosphate generation step between RINCAA and starvation-induced autophagy. Mechanistic analysis identified a ULK1-regulated phosphorylation region (K251-K269) located on PI4KB as a special event of RINCAA (Fig. 5), which is in line with several studies showing ULK1 as a major regulator of autophagy in response to different cancer signals in tumor and the potency of ULK1 inhibitors against tumor.^40, 119, 120^

Importantly, phosphorylation of the PI4K-Peptide-1 occurred in higher rates in RAS-mutated CRC and several cancer cell lines with the RAS mutation (Fig. 7a-c), suggesting that this phosphorylation event may be a special signature of cancer with a RAS mutation. Hence, PI4KB S256 and T263 phosphorylation may be a pathological form of non-canonical autophagy specific target for treating cancer with a RAS mutation. In support of this idea, blocking PI4KB S256 and T263 phosphorylation with a competing substrate peptide or overexpressing a phosphorylation-deficient PI4KB-SA (which likely competes with the function of endogenous phosphorylatable PI4KB) inhibits RINCAA and the growth of tumors with a KRAS mutation with equal or better effects compared to chloroquine (Fig. 6,8,9). While PI4KB plays a pivotal role in PI4P generation across various cellular processes, our findings indicate that S256 and T263 phosphorylation is not essential for PI4KB activity under normal conditions (Fig.5i). This suggests a potential for minimal side effects when targeting PI4KB S256 and T263 phosphorylation.

**Fig. 7.**
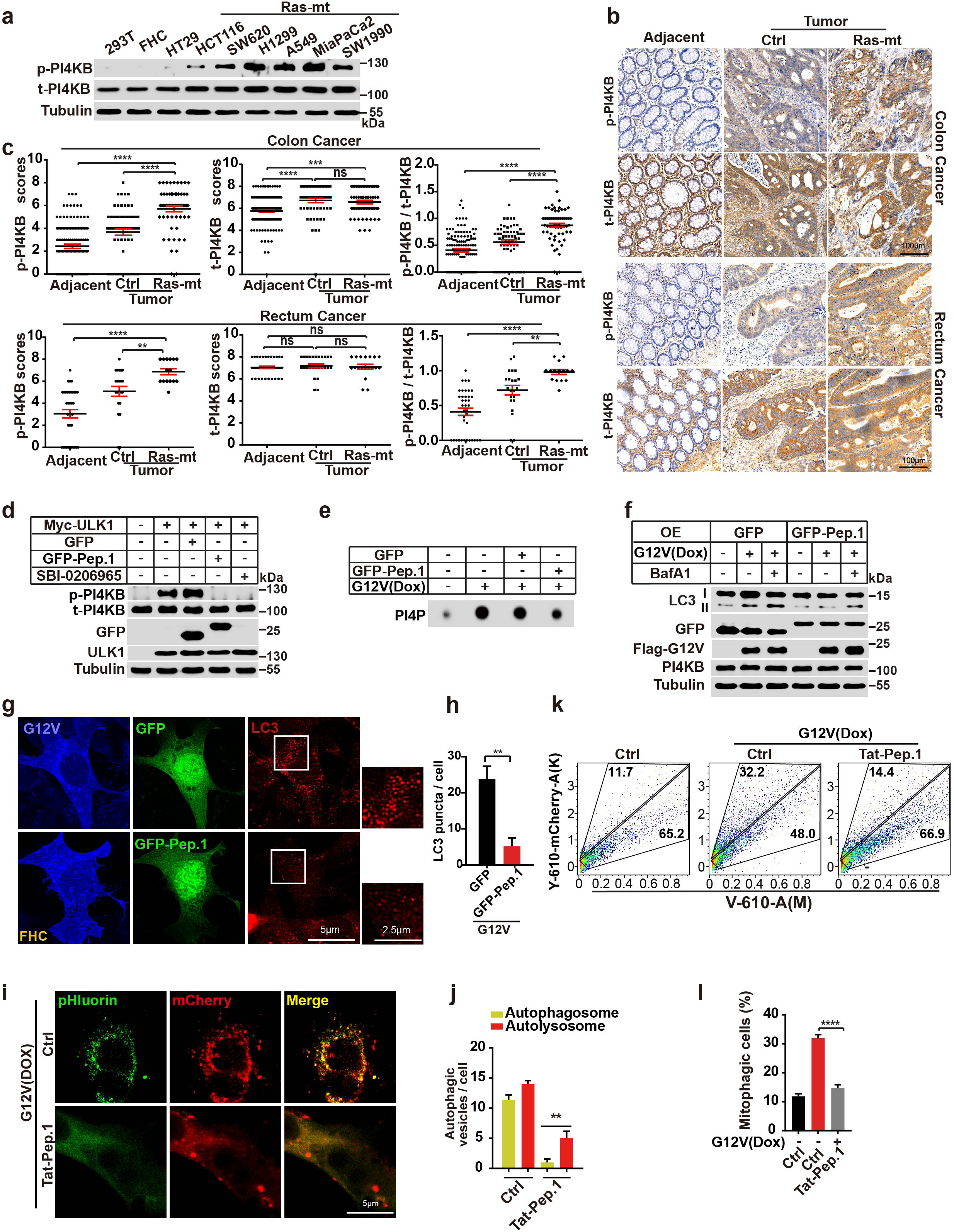
Targeting ULK1-mediated PI4KB phosphorylation on S256 and T263 could inhibit RINCAA. **a**. Immunoblot analysis of the level of p-PI4KB in different cancer cell lines. **b**. Representative immunohistochemical staining of p-PI4KB of the tissue microarrays containing CRC tissues (control and RAS-mutant) and adjacent normal tissues. **c**. Quantification of the p-PI4KB level of the tissue microarrays with colon cancer (Adjacent: n=119; Cancer: n=60 (Ctrl) and n=59 (RAS-mt)) and rectum cancer (Adjacent: n=37; Cancer: n=22 (Ctrl) and n=15 (RAS-mt)) (means ± SDs) patient samples. The statistical analysis was performed by the Wilcoxon signed rank test. **, p<0.01; ***, p<0.001; ****, p<0.0001. **d**. Immunoblot analysis of the PI4KB phosphorylation by the antibody against p-PI4KB in 293T cells expressing Myc-ULK1 with or without GFP-Peptide1 (GFP-Pep.1) overexpression or SBI-0206965 (10 μM) treatment. **d**. Dot blot analysis of the effect of GFP-Peptide1 on the KRAS(G12V)-induced PI4P generation. **e**. Immunoblot analysis of the KRAS(G12V)-induced LC3 lipidation when overexpressing GFP-Peptide1 (GFP-Pep.1) in the absence or presence of 500 nM Bafilomycin A1 for 1.5 h. **f**. Immunofluorescence analysis of the KRAS(G12V)-induced LC3 puncta with or without overexpressing GFP-Peptide1 (GFP-Pep.1). Representative images are shown. Scale bar sizes are indicated in the image. **g**. Quantification of the LC3 puncta in (g) (mean ±SEM). Three experiments (50 cells for each group/experiment) were performed for the statistics (two-tailed t-test). **, *p<*0.01. **h**. Detection of effect of GFP-Peptide1 (GFP-Pep.1) on autophagy flux by the tandem fluorescent LC3 system in KRAS(G12V) cells stably expressing mCherry-pHluorin- LC3B. Representative cell images are shown. Scale bar sizes are indicated in the image. **i**. Quantification of the yellow (RFP^+^GFP^+^) and Red (RFP^+^GFP^-^) LC3 puncta. Data are represented as mean ±SEM. Three experiments (50 cells for each group/experiment) were performed for the statistics (two-tailed t-test). **, *p<*0.01. **j**.FACS analysis of control and KRAS(G12V) cells co-expressing mt-Keima and Parkin with or without GFP-Peptide1 (GFP-Pep.1) expression using V610 and Y610-mCherry **k**. detectors (Beckman CytoFLEX LX). The FACS results are representative of at least three independent experiments. **l**. The percentage of cells with mitophagy based on Y610-mCherry/V610 calculated for (k). Data are represented as mean ±SEM. Three experiments were performed for the statistics (two-tailed t-test). ****, p<0.0001.

**Fig. 8.**
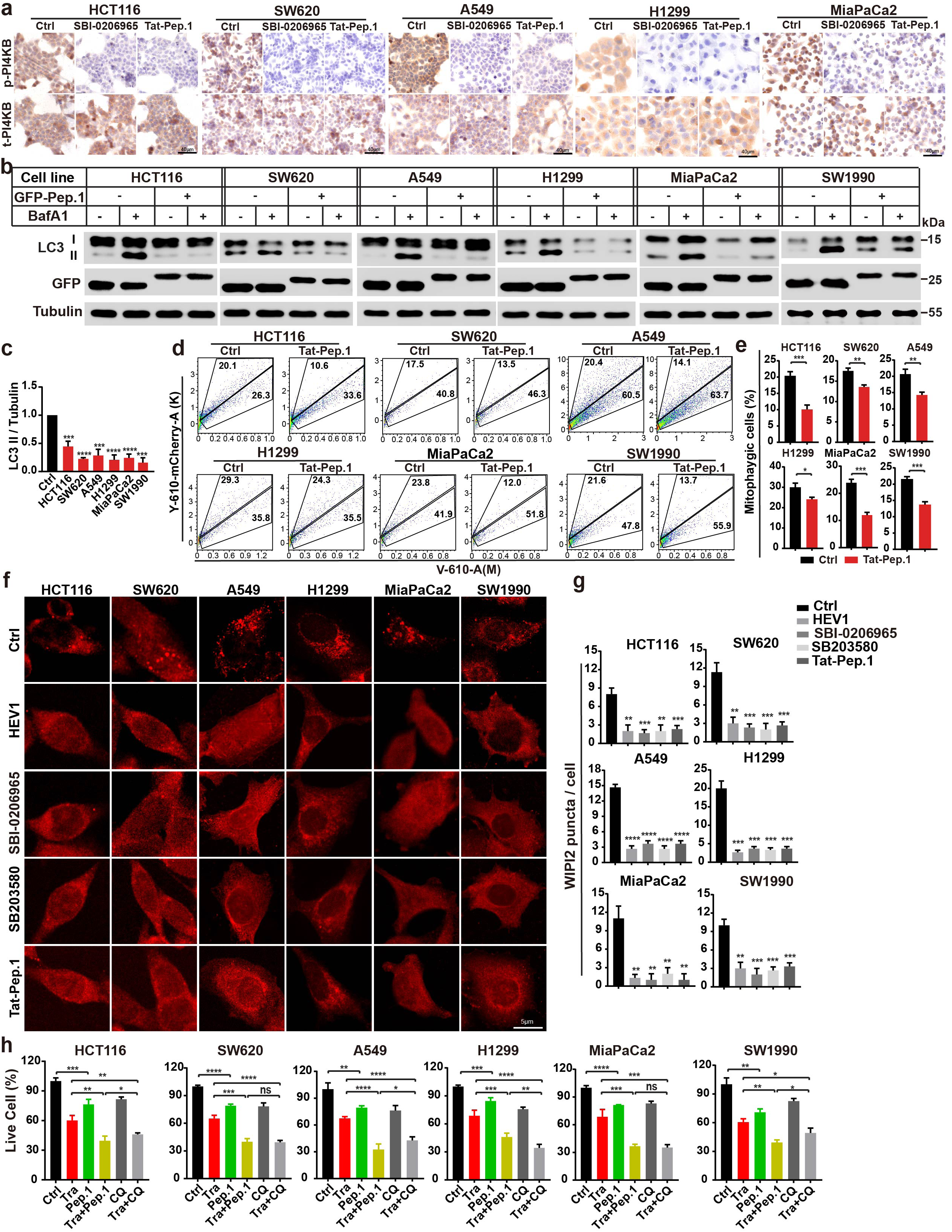
The PI4KB-Peptide1 inhibits RAS-mutant cell autophagy and growth. **a**. Immunohistochemical analysis of the sections of HCT116, SW620, A549, H1299 and MiaPaCa-2 cells treated with or without SBI-0206965 (10 μM) or Pepptide1 (Tat-Pep.1, 25 μM) for 2 h. Sections were stained with antibody against PI4KB and p-PI4KB as indicated. Scale bar sizes are indicated in the image. **b**. Immunoblot analysis of LC3 lipidation of HCT116, SW620, A549, H1299 and MiaPaCa-2 cells with or without overexpressing GFP-Peptide1 (GFP-Pep.1) in the absence or presence of 500 nM Bafilomycin A1 for 1.5 h. **c**. Quantification of the ratio of lipidated LC3 to tubulin with the control set as 1.00 (control cells with Bafilomycin A1) analyzed in (b) (mean ±SEM). Three experiments were performed for the statistics (two-tailed t-test). ***, p<0.01; ****, p<0.0001. **d**. FACS analysis of HCT116, SW620, A549, H1299 and MiaPaCa-2 cells co-expressing mt-Keima and Parkin with or without GFP-Peptide1 (GFP-Pep.1) expression using V610 and Y610-mCherry detectors (Beckman CytoFLEX LX). The FACS results are representative of at least three independent experiments. **e**. The percentage of cells with mitophagy based on Y610-mCherry/V610 calculated for (d). Data are represented as mean ±SEM. Three experiments were performed for the statistics (two-tailed t-test). ****, p<0.0001. **f**. Immunofluorescence analysis of the WIPI2 puncta of different cancer cell lines treated with HEV1 (10 μM), SBI-209626 (10 μM), SB203580 (10 μM) for 1.5 h or overexpressing GFP-Peptide1 (GFP-Pep.1). Representative cell images are shown. Scale bar sizes are indicated in the image. **g**. Quantification of the WIPI2 puncta in (g). Data are represented as mean ±SEM. Three experiments (50 cells for each group/experiment) were performed for the statistics (two-tailed t-test). **, p<0.01; ***, p<0.01; ****, p<0.0001. **h**. The CCK8 analysis of the cell proliferation of HCT116, SW620, A549, H1299 and MiaPaCa-2 cells treated with vehicle (Ctrl), Trametinib (Tra, 100 nM), Tat-Peptide1 (Pep.1, 5 μM), the combination of Trametinib and Tat-Peptide1 (Tra+Pep.1), chloroquine (CQ, 10 μM), the combination of Trametinib and chloroquine (Tra+CQ) for 96 h (mean ±SEM). Three experiments were performed for the statistics (two-tailed t-test). *, p<0.05; **, p<0.01; ***, p<0.001; ****, p<0.0001.

**Fig. 9.**
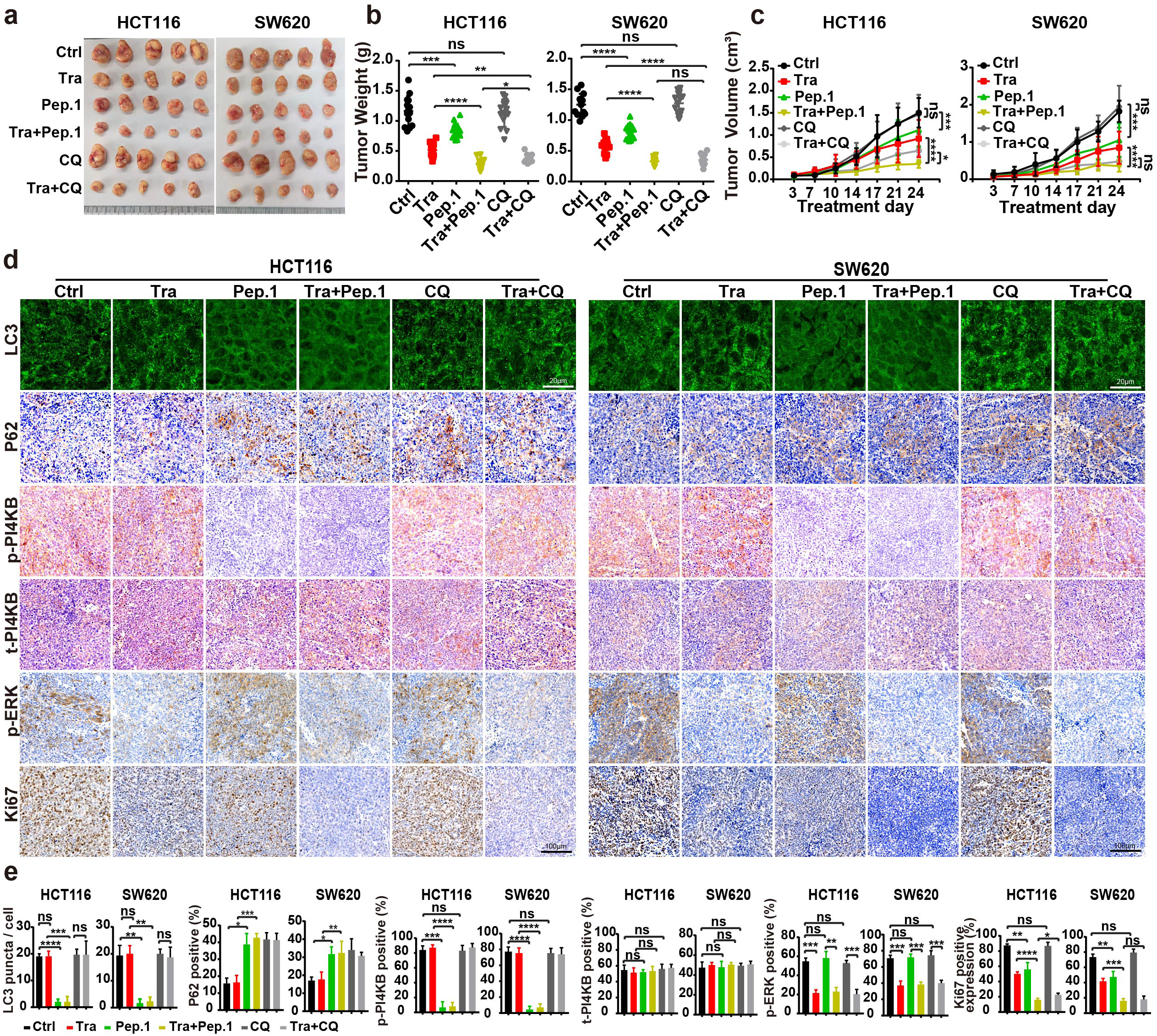
ULK1-mediated PI4KB phosphorylation on S256 and T263 may be a target for treating RAS-mutant cancer. **a**. Images of xenograft tumors of HCT116 and SW620 cells from mice treated with: (1) vehicle (Control); (2) Trametinib (Tra); (3) GFP-Peptide1 (GFP-Pep.1); or (4) the combination of Trametinib and GFP-Peptide1 (Tra+Pep.1) (5) chloroquine (CQ); (6) the combination of Trametinib and chloroquine (Tra+CQ) (n = 12 mice (one tumor/mice) in each group). **b**. Weights of the xenograft tumors in (j) (means ± SD). The statistical analysis was performed by two-tailed t-test. *, *p<*0.05; **, *p<* 0.01; ***, *p<*0.001. **c**. The growth curve of the xenograft tumors in (j) (means ±SD). Statistical analysis was performed by two-way-ANOVA; *, *p<*0.05; **, *p<* 0.01; ***, *p<*0.001; ****, *p<*0.0001. **d**. Immunofluorescence and immunohistochemical analysis of sections of the xenograft tumors in (j). Sections were stained with antibody against LC3, P62, p-PI4KB, PI4KB, p-ERK1/2 or Ki67, as indicated. Scale bars are located at the bottom right of the images. **e**.Statistical analysis of the numbers of LC3 puncta, P62 positive rates, p-PI4KB positive rates, t-PI4KB positive rates, p-ERK positive rates, and levels of Ki67 expression in (m) (means ±SD). Statistical analysis was performed by two-tailed t-test. *, *p<*0.05; **, *p<*0.01; ***, *p<*0.001; ****, *p<*0.0001.

**Fig. 10.**
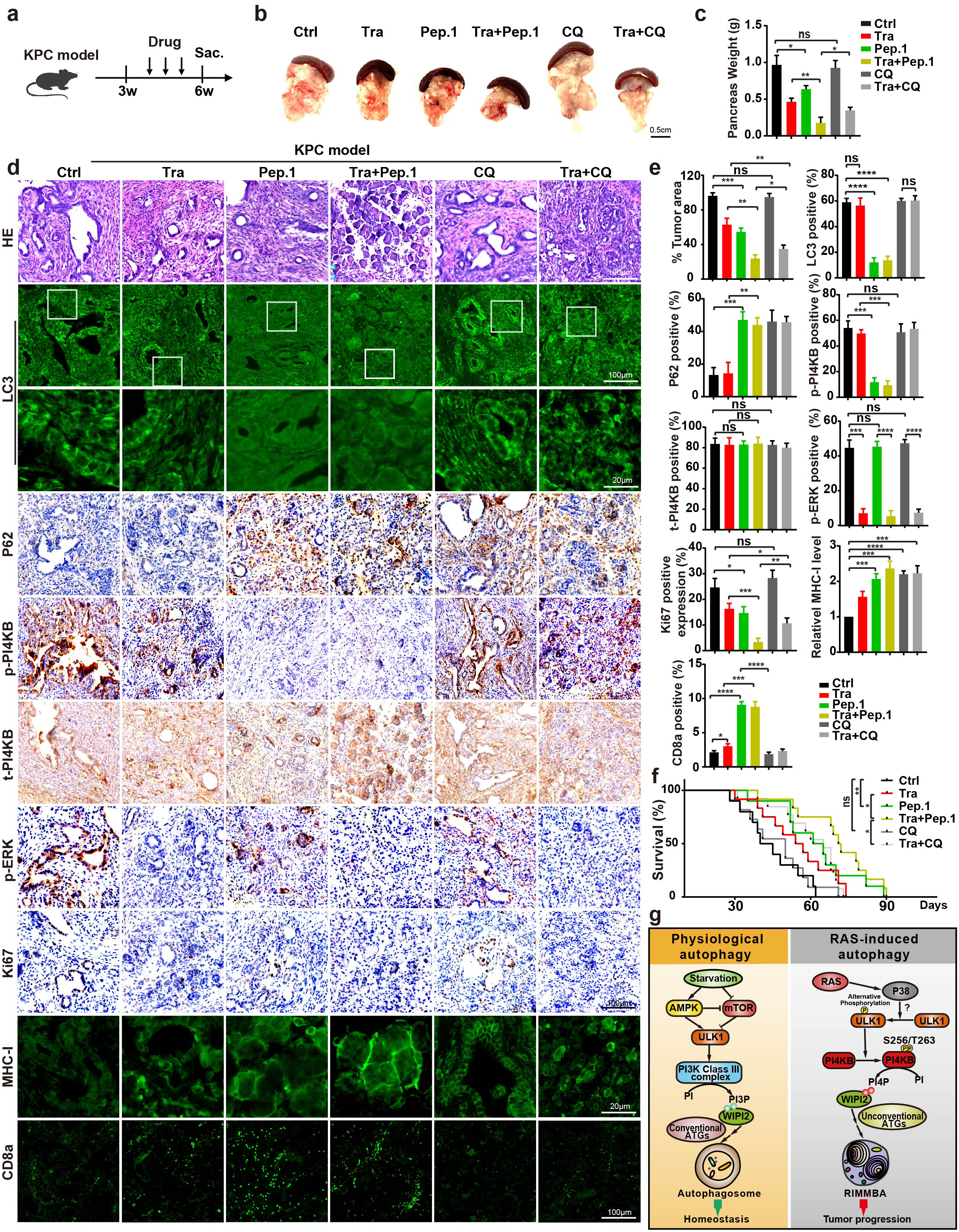
Inhibition of tumors by PI4KB-peptide 1 in a mice KPC model of pancreatic cancer. **a**. Schematic of experimental design in KPC mice. **b**. Representative images from primary pancreatic tumors at endpoint analysis in indicated treatment groups: vehicle (Ctrl); Trametinib (Tra); Tat-Peptide1 (Pep.1); the combination of Trametinib and Tat-Peptide1 (Tra+Pep.1); chloroquine (CQ); the combination of Trametinib and chloroquine (Tra+CQ). **c**. Weights of primary pancreatic tumors in (b) (means ±SD) (n = 10 mice). The statistical analysis was performed by two-tailed t-test. *, p<0.05; **, p< 0.01. **d**. Immunofluorescence and immunohistochemical analysis of sections of the xenograft tumors in (b). H&E staining is provided to show the state of the tissue in each condition. Sections were stained with antibody against LC3, P62, p-PI4KB, PI4KB, p-ERK1/2 or Ki67, as indicated. Scale bars are located at the bottom right of the images. **e**. Statistical analysis of tumor area and the numbers of LC3 puncta, P62 positive rates, p-PI4KB positive rates, t-PI4KB positive rates, p-ERK positive rates, and levels of Ki67 expression in (d) (means ±SD). Statistical analysis was performed by two-tailed t-test. *, p<0.05; **, p<0.01; ***, p<0.001; ****, p<0.0001. **f**. Kaplan–Meier curves of KPC mice were shown. The mice were treated with vehicle (Ctrl, n = 10 mice); Trametinib (Tra, n = 11 mice); Tat-Peptide1 (Pep.1, n = 10 mice); the combination of Trametinib and Tat-Peptide1 (Tra+Pep.1, n = 12 mice); chloroquine (CQ, n = 11 mice); the combination of Trametinib and chloroquine (Tra+CQ, n = 13 mice). **g**. A model for oncogenic RINCCA.

In addition to promoting autophagy, RAS mutations have been reported to increase macropinocytosis, facilitating enhanced nutrient uptake in cancer cells.^121–123^ Although chloroquine has been shown to inhibit both autophagy and macropinocytosis, PI4KB depletion similarly inhibits both processes (Fig.2, and Supplementary information, Fig.

S3,S6i, j). However, the more specific RINCAA inhibitor, PI4KB-Peptide-1, selectively blocks autophagy without affecting macropinocytosis (Supplementary information, Fig. S6k-n). Notably, despite this selectivity, PI4KB-Peptide-1 demonstrated superior antitumor efficacy compared to chloroquine, suggesting that preserving immune function and disrupting tumor metabolic remodeling are more critical for effectively treating RAS- mutant cancers than simply reducing nutrient scavenging. The high specificity of short peptides like PI4KB-Peptide-1 in targeting RINCAA within RAS-mutant cancers, while maintaining immune activity, underscores their potential as a promising anticancer approach. Future research should focus on developing small-molecule inhibitors targeting PI4KB phosphorylation at S256 and T263 and assessing their clinical effectiveness for enhanced therapeutic outcomes.

## ACKNOWLEDGMENTS

Dr. Liang Ge and Dr. Min Zhang are deeply grateful for the postdoc training done in Dr. Randy Schekman’s lab at University of California, Berkeley. We thank Dr. Hong Zhang (Institute of Biophysics, Chinese Academy of Sciences, China) and Dr. Yueguang Rong (Huazhong University of Science and Technology, China) for plasmids. We thank Dr. Charles J. David and Dr. Mo Chen (Tsinghua University) for the KPC mice and insightful discussions on the data. We thank Dr. Wei Liu (Zhejiang University) and Dr. Li Yu (Tsinghua University) for helpful suggestions on the study. We thank Ningjia Ma, Lingzhu Zhang, and Zinan Liu for technique assistance. We would like to thank Shuoguo Li, Yun Feng, Tongxin Niu and Xiaoyun Zhang from Center for Biological Imaging (CBI), Institute of Biophysics, Chinese Academy of Science for their helps of taking and processing cryo- CLEM / ET freeze fracture work. The authors would like to acknowledge the assistance of Imaging Core Facility, Technology Center for Protein Sciences, Tsinghua University. The work is funded by National Natural Science Foundation of China (32225013, 32130023, 92254302, 32370728), National Key R&D Program of China (2019YFA0508602; 2021YFA0804802), Tsinghua University Dushi Program, China Postdoctoral Science Foundation (BX20240186 and 2024M761616), Shuimu Tsinghua Scholar Program.

## AUTHOR CONTRIBUTIONS

1. X. W. and S. Li. designed experiments, performed experiments, analyzed data, and wrote the manuscript. S. Lin., T. Z., Z.H., Y. L., and J. W. performed experiments. Y. H, and H.
2. D. analyzed data. M. Z., D. F. and L. G. designed experiments, analyzed data, and wrote the manuscript.

