## supplementary file for "Oncogenic RAS Induces a Distinctive Form of Non-Canonical Autophagy Mediated by the P38-ULK1-PI4KB Axis"

Xiaojuan Wang *et al.*

Corresponding authors:

Min Zhang:

Du Feng:

Liang Ge:

**The PDF file includes:**

Materials and methods

Supplementary figures 1-6

Supplementary Tables 1-3

**Other Supplementary Material for this manuscript includes the following:**

Supplementary Movie 1

### MATERIALS AND METHODS

#### Cells

Cells were maintained in Dulbecco's modified Eagle's medium (DMEM) (HEK293T, HCT116, SW620, A549, H1299, PANC1, MiaPaCa2, SW1990, MEF), or RPMI-1640 (FHC) with 10% FBS at 37°C in 5% $CO_2$. HEK293T cells stably expressing Flag-KRAS(G12V) (Flag-KRAS(G12V)-293T stable) were obtained by lentiviral infection followed by selection with antibiotics. For lentiviral transduction, HEK293T cells were transfected with plasmids (pMD2.G and psPAX2). Viruses were harvested at 60–72 h post-transfection. And the viral supernatant was centrifuged at 600× g for 5 min to remove cell debris. The indicated cells were infected with the viral supernatant diluted in fresh medium (30% viral supernatant) with 10 μg/mL polybrene, which was exchanged to growth medium after 24 h. Transduced cells were selected and cultured in DMEM supplemented with 10% FBS, 150 μg/mL Zeocin (Sigma), and 15 μg/mL Blasticidin (Sigma). Target-gene expression was confirmed via SDS-PAGE and immunoblot.
Primary antibodies used in this study are listed in Supplementary information, Table S2.

#### Mice

The mice experiments were approved by the Institutional Animal Care and Use Committees at Tsinghua University.

For xenograft studies, nude mice were purchased from Charles River (Beijing) and housed in ventilated cages in a temperature and light-regulated room in an SPF facility and received food and water ad libitum. Xenografted tumors were established by sub-cutaneous injection upper thigh of $2 \times 10^6$ HCT116 or SW620 cells resuspended in 100 μL of Matrigel (Yeasen, 40183ES10) into the flanks of 4-week-old male NOD/SCID mice and allowed to establish. Treatment was then initiated with vehicle control (corn oil), trametinib at 1 mg/kg, Tat-Peptide1 (Tat -Pep.1) at 40mg/kg, chloroquine at 50 mg/kg, the combination of trametinib plus Tat -Pep.1 at the aforementioned dosages, or the combination of trametinib plus chloroquine at the aforementioned dosages via intraperitoneal injection twice a week. Tumors were measured twice weekly via calipers and tumor volume was calculated by volume = $4/3 \times \pi \times (((\text{length} + \text{width})/2)/2)^3$. More than 12 tumors per group were analyzed. The tumors were removed, photographed, and weighed, and the average weights of the tumors were calculated. Significance of difference in tumor size was calculated by a two-tailed t-test.

LSL-*Kras*$^{G12D}$, *p53*$^{F/F}$, *Pdx1-Cre* mice (KPC) were a gift from Charles J. David (Tsinghua University). Age- and sex-matched (except where indicated otherwise) male and female mice of the genotype were generated as littermates for use in experiments. Treatment was then initiated with vehicle control (corn oil), trametinib at 1 mg/kg, Tat-Peptide1 (Tat - Pep.1) at 40mg/kg, the combination of trametinib plus Tat -Pep.1 at the aforementioned dosages, chloroquine at 40 mg/kg or the combination of trametinib plus chloroquine at the aforementioned dosages via intraperitoneal injection three times a week.

**Plasmids, siRNA oligos, and transfection**

Human KRAS(G12V), KRAS(G12D), or KRAS(G12C) was cloned into pLenti-CMV vector. The mCherry-pHluorin-LC3B, mCherry-ATG4B-C74S, OSBP-PH, WIPI2-GST Flag-ULK1, Myc-ULK1, HA-ULK1, EGFP-PI4KB, HA-PI4KB and GFP-Pep.1 plasmids were generated by PCR and ligation. HA-PI4KB-SA, HA-PI4KB-D656 and Myc-ULK1-K46I plasmids were generated by site mutagenesis PCR. The siRNAs and shRNAs are listed in Supplementary information, Table S3. Transfection of PI4P (Avanti Polar Lipids) was performed using Unlabeled Shuttle PIP Carrier 3 (Echelon Biosciences). Transfection of plasmids was performed using PEI (Polysciences, Inc.) for HEK293T and X-tremeGENE HP (Roche) for FHC according to the manufacturers’ protocols. The siRNA transfection was performed with Lipofectamine RNAiMAX (Invitrogen) according to the manufacturer’s protocol.

**Reagents**

Doxycycline Hyclate was purchased from Beyotime (Cat# ST039B). Inhibitors including Bafilomycin, Wortmannin, AMG510, SAR405, PIK-III, VPS34-IN1, PIK93, SB203580, JNK-IN-8, FR180204, MK2206, RBC8, SBI-0206965 were purchased from Selleck (Cat# S1413, S2578, S8830, S7682, S7683, S7980, S1489, S1076, S4901, S7524, S1078, S7606, and S7885, respectively). T-00127-HEV1 was purchased from MedChemExpress (Cat# HY-108313). YM201636 was purchased from Topscience, USA (Cat# T6110). PI4P was purchased from Echelon Biosciences (Cat# p4016-2). Digitonin was purchased from BIOSYNTH (Cat# D-3200). CIP was purchased from Sigma (Cat# P4978).

**Peptide synthesis**

L-amino acid peptides were synthesized by Scilight Biotechnology LLC. The Tat-peptide1 (Tat-Pep.1) peptide sequence, YGRKKRRQRRRGGELPSLSPAPDTGLSPSK, consisted of 11 amino acids from the Tat PTD at the N-terminus, a GG linker to increase flexibility, and at the C-terminus, 17 amino acids derived from PI4KB (253-269). The control peptide (Tat-scrambled) consisted of the Tat protein transduction domain, a GG linker, and a scramble sequence (YGRKKRRQRRRGGVGNDFFINHETTGFATEW).

**Immunofluorescence**

The cells were incubated with 4% cold paraformaldehyde for 20 min at room temperature. The cells were further permeabilized with 0.1% Triton X-100 diluted in PBS at room temperature for 10 min, followed by blocking with 10% FBS diluted in PBS for 1 h and the primary antibody incubation for 1 h at room temperature. Cells were washed three times with PBS, followed by the secondary antibody incubation for 1 h at room temperature.

For the tissue immunofluorescence, 5 μm sections of the paraffin-embedded tumors were kept at 60°C for 24 h in the oven followed by deparaffinized with xylene and hydrated with an ethanol gradient (100-70%). After successively incubating with antigen retrieval solution (ZSGB Biotechnology Company; Beijing, China) and 3% $H_2O_2$ for 30 min, the slides were rinsed with water and incubated with the primary antibody overnight at 4°C. The next day, the slides were rinsed and incubated with the corresponding secondary antibody for 1 h at room temperature.

Fluorescence images were acquired using the Olympus FV3000 confocal microscope. Quantification was performed using ImageJ.

**HaloTag-LC3B processing assay**

HEK293T cells stably expressing the Tet-on KRAS(G12V) system and HaloTag-LC3B were used for autophagic flux analysis. After 36 hours of induction with doxycycline (final concentration 1 µg/ml), the cells were pulse-labeled for 20 minutes with 100 nM tetramethyl rhodamine (TMR)-conjugated ligand (Promega, G8251) in nutrient-rich medium. Following two washes with phosphate-buffered saline (PBS), the cells were further incubated for 2 hours in fresh medium containing Dox (final concentration 1 µg/ml) alone or in combination with 100 nM Bafilomycin A1 (Baf A1). After incubation, cells were lysed and 20 µg of protein per sample was subjected to SDS-PAGE. For in-gel fluorescence imaging, the gel was immediately imaged using ChemiDoc Imaging System (Bio-Rad) after SDS-PAGE. For immunoblotting, proteins were transferred from the SDS-PAGE gel to Immobilon-P polyvinylidene difluoride (PVDF) membranes (Millipore, IPVH00010). Following incubation with the appropriate antibodies, the signals were detected using chemiluminescent HRP substrate and ChemiDoc Imaging System (Bio-Rad). Band intensities were quantified using the Gel Analyzer tool in the open-source image processing software Fiji.

**Co-immunoprecipitation (co-IP) and immunoblot**

For co-IP, the cells were lysed on ice for 30 min in the IP buffer (50 mM Tris/HCl, pH 7.4; 150 mM NaCl; 1 mM EDTA; 0.5% NP40; 10% glycerol) with protease inhibitor mixture, and the lysates were cleared by centrifugation. The resulting supernatants were incubated with indicated antibody-conjugated agarose beads and rotated for 3 h at 4°C. Then the agaroses were washed five times with the IP buffer and checked by immunoblot.

### Immunoprecipitation and in vitro kinase assay

Cells expressing indicated proteins were lysed in ice-cold lysis buffer (20 mM Tris, pH 7.5; 150 mM NaCl; 0.3% [vol/vol] Triton X-100; 5 mM EDTA) supplemented with Complete EDTA-free protease inhibitor cocktail (Roche) and PhosStop phosphatase inhibitor cocktail (Roche). The lysates were centrifuged, and the supernatant was incubated with indicated antibody-conjugated agarose beads at 4°C for one hour. Following incubation, the beads were washed three times with the lysis buffer (above-mentioned) and then once with kinase reaction buffer (20 mM HEPES, pH 7.5; 20 mM $MgCl_2$; 25 mM beta-glycerophosphate; 2 mM dithiothreitol; 100 µM sodium orthovanadate). Subsequently, the beads were incubated in a final volume of 20 µl reaction buffer containing 20 µM ATP and 5 µg purified GFP-PI4KB protein at 30°C for 15 minutes. The reactions were stopped by adding 5× SDS sample buffer and heating to 65°C for 5 minutes. Proteins were then transferred to PVDF membranes for analysis by immunoblotting.

### Dot blot

For determination of PI4P, total lipids were extracted from HEK293T cells. Each 300 mL cell suspension was incubated with 1.2 mL chloroform/methanol solution (chloroform: methanol = 1:2) and vortexed for 30 s and shaken for 1 h at 180 rpm at 37℃. The chloroform phase was collected and evaporated by a stream of nitrogen gas over the lipid solution and further dried in an incubator at 37℃ for 1 h. Dried lipids were suspended in absolute ethyl alcohol. The lipids solution was measured and used as a standard (PC) to normalize lipids concentration. The lipids were applied to the nitrocellulose membrane (Millipore; Bedford, MA) and left to dry for 1 h. The membrane was incubated with the PI4P antibody overnight at room temperature. The next day, the membrane was washed and incubated with an anti-mouse antibody (diluted 1:1000 in the blocking solution) for 1

h. Finally, the PI4P level recognized by the antibody was revealed by treatment with chemiluminescent.

For determination of WIPI2 binding to lipids, purified WIPI2 was used to overlay membrane-immobilized phospholipid membranes (PIP Strips, Echelon, Cat#P-6001). Anti-WIPI2 and anti-mouse secondary antibodies were used to detect the proteins by scanning with the ChemiDoc Imaging Systems.

### qRT-PCR

Total RNA was isolated from different cell lines using TRIzol reagent (Beyotime, Cat No. R0016) according to the manufacturer's instruction. Equal amounts of RNA were reverse transcribed into cDNA using a Revert Aid First Strand cDNA synthesis kit (Abclonal, E047-01B) according to the manufacturer's instruction. Quantitative PCR was performed using an ABI Step One Plus system. The PCR reactions were carried out in 10 μL reactions using SYBR Green PCR master mix (Abclonal, RK20429) and 0.5 μM specific primers. The primers used for PCR are shown in Supplementary Table 3.

### Electron microscopy (EM) and APEX2-DAB staining

For DAB staining and preparation of cultured cells for EM, 293T-KRAS(G12V) cells were transfected with APEX2-LC3B and concurrently induced with doxycycline (final concentration 1 μg/ml). After 36 hours of induction, the cells were fixed using 2% glutaraldehyde in buffer (100 mM sodium cacodylate with 2 mM CaCl2, pH 7.4), then quickly moved to ice. Cells were kept between 0 and 4 °C for all subsequent steps until resin infiltration. After 30–60 min, cells were rinsed 5× 2 min in chilled buffer, treated for 5 min in buffer containing 20 mM glycine to quench unreacted glutaraldehyde followed by 5 × 2 min rinses in chilled buffer. A freshly diluted solution of 0.5 mg/mL (1.4 mM) DAB tetrahydrochloride or the DAB free base (Sigma) dissolved in HCl was combined with 0.03% (v/v) (10 mM) H2O2 in chilled buffer, and the solution was added to cells for 5 min. The generation of reaction product could be monitored by transmitted LM. To halt the reaction, the DAB solution was removed, and cells were rinsed 5× 5 min with chilled buffer. Post-fixation staining was performed with 2% osmium tetroxide for 30 min in chilled buffer. Cells were rinsed 5× 2 min in chilled distilled water and placed in chilled 2% aqueous uranyl acetate overnight. The samples were then dehydrated in a cold-graded ethanol series (20%, 50%, 70%, 90%, 100%, 100%) 2 min each, rinsed once in room temperature anhydrous ethanol to avoid condensation, and infiltrated in Durcupan ACM resin (Electron Microscopy Sciences) using 1:1 (v/v) anhydrous ethanol and resin for 30 min, and 100% resin 2× 1 h. Finally, the samples were embedded into fresh resin and polymerized in a vacuum oven at 60 °C for 48 h. DAB-stained areas of embedded cultured cells were identified by transmitted light, and the areas of interest were sawed out using a jeweler's saw and mounted on dummy acrylic blocks with cyanoacrylic adhesive (Krazy Glue, Elmer's Products). The coverslip was carefully removed, the block trimmed, and ultrathin (80 nm thick) sections were cut using an ultramicrotome (Leica Ultracut UTC6). Samples were imaged under the HT-7800 120kv transmission electron microscope.

**In situ cryo-correlative light and electron microscopy (CLEM) / electron tomography (ET)**

For sample preparation, FHC cells expressing BFP-KRas-G12V, EGFP tagged OSBP-PH and mCherry-LC3B were seeded onto ultraviolet-sterilized ELI grids (T11012ss-ELI, TIANLD) and cultured in complete DMEM supplemented with 10% FBS and 1% penicillin and streptomycin. After 48 h of culture with 5% CO2 at 37 °C, the grids were subjected to plunge freezing by backside blotting and vitrification using a Leica EM GP (Leica Microsystems). Next, the cryo-vitrified grid was assembled into a FEI AutoGrid (Thermo Fisher Scientific, USA) for the subsequent cryo-transfer and imaging.

For the HOPE-SIM screening, Cryo-SIM images were acquired on the HOPE-SIM imaging system with a ×100 0.9 dry objective (Nikon), a laser combiner of three lasers with wavelengths of 405 nm (50 mW, OBIS 405LX, Coherent), 488 nm (200 mW, SAPPHIRE 488-200 CW, Coherent), and 561 nm (200 mW, SAPPHIRE 561-200 CW, Coherent) and a high-sensitivity sCMOS camera (Prime 95B, Photometrics). Each raw image has a pixel size of 120 nm and a field of view (FOV) of 1,200 × 1,200 pixels. Then, the final raw image stack has a voxel size of 120 × 120 × 250 nm.

For cryo-FIB milling, cryo-lamellae were prepared by cryo-FIB milling using the FEI Helios NanoLab 600i (Thermo Fisher Scientific, USA). Cryo-FIB milling was accurately navigated by monitoring the real-time fluorescence signal of the target molecule by using ELI trifocal microscope. CLEM correlation operations were performed in 3D-View software.

For cryo-ET data collection, the cryo-lamellae were loaded into a FEI Titan Krios G3i (Thermo Fisher Scientific, USA) that was equipped with a Gatan GIF K2 4k × 4k camera (Gatan, Inc., Warrendale, PA) and operated at 300 kV in low-dose mode. Tilt series were collected bidirectionally with SerialEM software using a tilt range of -55° to 50° in 2° intervals, nominal magnification of 33,000×, defocus of −6 μm. All procedures for cryo-EM imaging were performed under low-dose conditions.

For data processing, image movies were processed by motion correction and CTF estimation in Warp. The produced tilt series were aligned using IMOD.[142] Amira 10.0 (Thermo Fisher Scientific, USA) was applied to segment structures of the cryo-ET images. The characteristics of the structures were labeled manually and colored differently. All movies were generated in Imaris 10.0 (Oxford Instruments).

**Mass spectrometry (MS) and Phos-Tag gel analysis**

293T cells expressing HA-PI4KB with or without Myc-ULK1 were cultured on three 10-cm dishes to 100% confluency. The small-molecule kinase inhibitor SBI-0206965 was added to the culture to inhibit the ULK1. Then agarose coupled with anti-HA antibodies was employed to immunoisolate the HA-PI4KB and eluted with the HA peptide. Purified HA-PI4KB was analyzed by SDS–PAGE and visualized by Coomassie Brilliant Blue staining. Specific HA-PI4KB bands were subjected to mass spectrometry analysis to identify post-translational modifications including phosphorylation. Mass spectrometry was performed by the Protein Chemistry and Proteomics Center at Tsinghua University. Phos-Tag gel analysis (Wako, 199-17391) was applied to confirm the phosphorylation of PI4KB according to the manufacturer's instruction.

**Liposome floatation and pelleting assay**

POPC (1-palmitoyl-2-oleoyl-glycero-3-phosphocholine), POPS (1-palmitoyl-2-oleoyl-sn-glycero-3-phospho-L-serine), DOPE (1,2-dioleoyl-sn-glycero-3-phosphoethanolamine), cholesterol, PI3P, PI4P, and PI5P were purchased from Avanti Polar Lipids. Lipids were mixed with a ratio of 4:2.5:2.5:10 for POPC: POPS: DOPE: cholesterol plus 1% PI3P, PI4P, or PI5P. The preparation of small unilamellar vesicles (SUVs) was described previously(*142*), with the lipids mixture above. The lipids mixtures were dried with a nitrogen stream and further dried for 1 h at 37°C. The lipid film was then hydrated sufficiently hydrated completely with the HEPES buffer (20 mM HEPES, pH 7.4; 150 mM NaCl) and subjected to 10 cycles of freezing in the liquid nitrogen and thawing in a 42°C water bath. Finally, the liposomes were extruded 20 times through the polycarbonate film with a specific pore size to generate the SUVs.

For the floatation assay, purified WIPI2 was added into 120 μL SUVs solution with the described concentration and incubated for 30 min at room temperature. In order to remove the free proteins, a membrane floatation procedure was performed as described before(*142*). Briefly, 480 μL 50% OptiPrep was added to the 120 μL solution above. The mixture was overlaid successively with 480 μL 30% OptiPrep and 90 μL HEPES buffer, then centrifuged at 100,000xg for 2 h. The 60 μL top fraction was collected. 1 μL of the top fraction was taken to measure the PC concentration with a microplate spectrophotometer. The rest was added with 5 × SDS loading buffer and analyzed by immunoblot.

**Cell proliferation assay**

Cell proliferation was assessed using a CCK-8 assay kit (Dojindo Molecular Technologies). Briefly, cells were seeded in 96-well plates. 10 μL CCK-8 solution was added to each well containing 100 μL culture medium and incubated for 2 h at 37°C. The absorbance was measured at 450 nm wavelength using an ELISA plate reader. For the cell proliferation assays, cell growth was analyzed on the first and the fourth day.

**Immunohistochemical (IHC) staining**

Pancreata or tumors were dissected and fixed in 4% paraformaldehyde in PBS and embedded in paraffin. 5 μm sections were prepared and stained with hematoxylin and eosin (H&E). For IHC, 5 μm sections of the paraffin-embedded tumors or pancreata were kept at 60°C for 24 h in the oven followed by deparaffinized with xylene and hydrated with an ethanol gradient (100-70%). After successively incubating with antigen retrieval solution (ZSGB Biotechnology Company; Beijing, China) and 3% $H_2O_2$ for 30 min, the slides were rinsed with water and incubated with the primary antibody overnight at 4°C. The next day, the slides were rinsed and incubated with the corresponding secondary antibody (ZSGB Biotechnology Company; Beijing, China) for 30 min followed by 3,3′-diaminobenzidine (DAB) and hematoxylin staining, respectively.

Tissue microarrays containing tumor tissues and their corresponding adjacent normal tissues from colon cancer and rectum cancer were obtained from Shanghai Wellbio Technology Co., Ltd. The samples were paraffin-embedded. Patient consent and approval from the Institutional Research Ethics Committee were obtained for the usage of these clinical materials for research purposes. Paraffin-embedded tissue sections (4 mm) were prepared according to standard methods, and the expression of p-PI4KB (1:1000 dilution) was detected by IHC staining. The slides were assessed by pathologists who were blinded to the experimental results and patient outcomes.

### IHC scoring

A modified labeling score (H score) which is calculated by using percentage of positive stained cancer cells and their intensity per tissue core. The intensity of stain was evaluated by an immunostaining score, which was calculated as the sum of the proportion and intensity of the stained tumor cells. Briefly, a proportion score, which represented the estimated proportion of positively stained tumor cells (0, none; 1, <1/100; 2, >1/100 to <1/10; 3, >1/10 to <1/3; 4, >1/3 to <2/3, and 5, >2/3.), was obtained. Next, an intensity score, which indicated the average intensity of positively stained tumor cells (0, none; 1, weak; 2, intermediate; and 3, strong), was obtained. The proportion and intensity scores were then added to obtain a total score, which ranged from 0 to 8.

### Statistical analysis

The ways of quantification of each experiment have been provided in the Method Details. The statistical information of each experiment, including the statistical methods, the P values and numbers (n), were shown in the figures and corresponding legends. Statistical analyses were performed with GraphPad Prism.

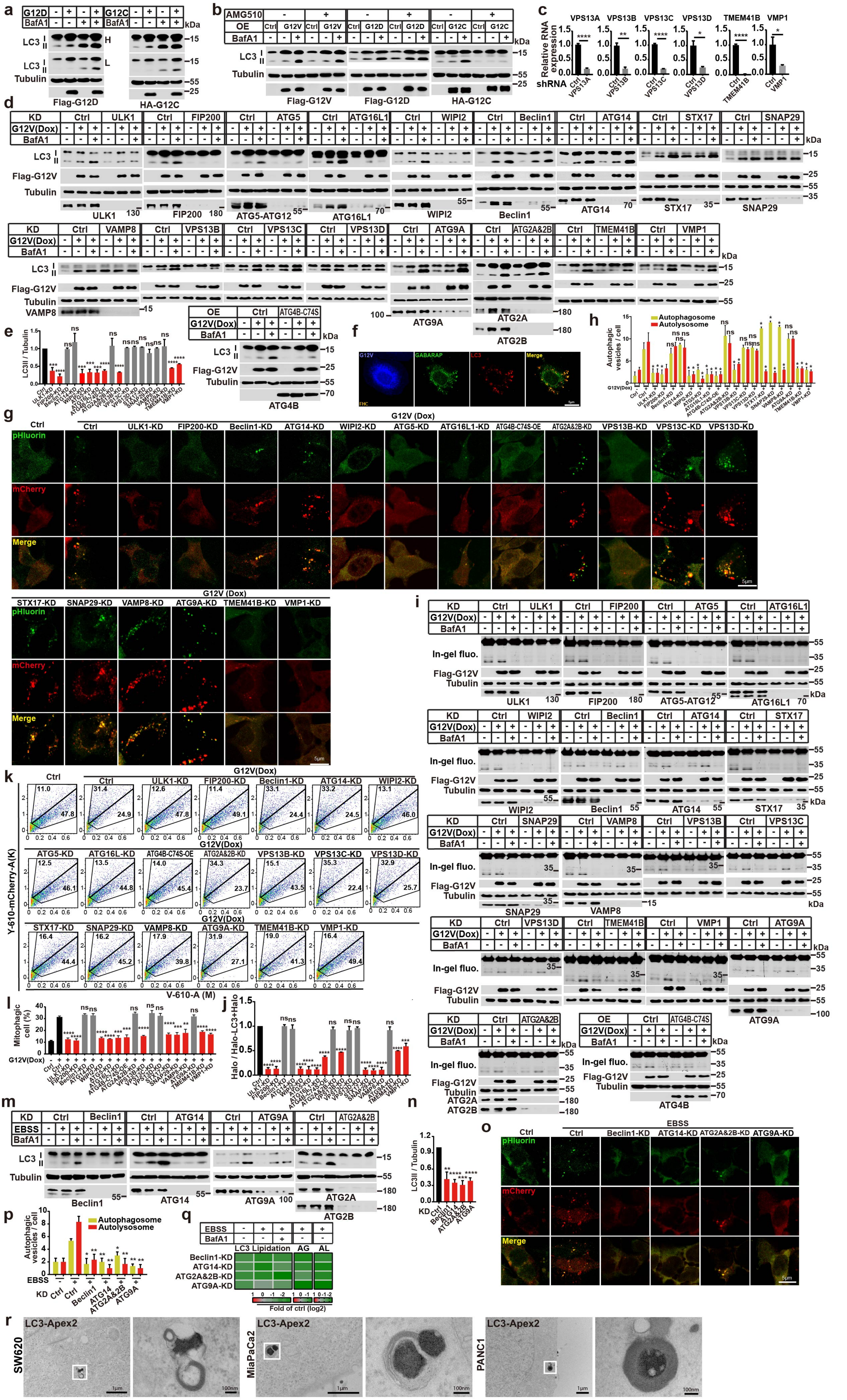
a
b
c
d
e
f
g
h
i
j
k
l
m
n
o
p
q
r
Flag-G12D
HA-G12C
Flag-G12V
LC3 I
II
Tubulin
BafA1
AMG510
OE
KD
G12V(Dox)
EBSS
Relative RNA expression
shRNA
VPS13A
VPS13B
VPS13C
VPS13D
TMEM41B
VMP1
ULK1
FIP200
ATG5-ATG12
ATG16L1
WIPI2
Beclin1
ATG14
STX17
SNAP29
VAMP8
ATG9A
ATG2A
ATG2B
ATG4B
ATG4B-C74S
LC3II / Tubulin
Autophagic vesicles / cell
Autophagosome
Autolysosome
pHluorin
mCherry
Merge
G12V (Dox)
In-gel fluo.
Y-610-mCherry-A(K)
V-610-A (M)
Mitophagic cell (%)
Halo / Halo-LC3+Halo
LC3 Lipidation
AG
AL
Fold of ctrl (log2)
LC3-Apex2
SW620
MiaPaCa2
PANC1
kDa

**Figure. S1 Characterization of KRAS(G12V)-induced autophagy**

**a.** Immunoblot analysis of the LC3 lipidation of the cell lysates from the control cells and the cells transfected with KRAS(G12D) or KRAS(G12C) plasmids.

**b.** Immunoblot analysis of the LC3 lipidation of the cell lysates from the control or KRAS(G12V), KRAS(G12D), KRAS(G12C) cells treated with 10 μM AMG510 for 1.5 h in the absence or presence of 500 nM Bafilomycin A1 for 1.5 h.

**c.** Relative RNA expression of cells transfected with control or shRNAs against VPS13B, VPS13C, VPS13D, TMEM41B and VMP1 (mean ± SEM). Three experiments were performed for the statistics (two-tailed t-test). *, $p<0.05$; **, $p<0.01$; ***, $p<0.001$; ****, $p<0.0001$.

**d.** Immunoblot analysis of the LC3 lipidation from KRAS(G12V)-cells with control or knocking down of ULK1, FIP200, ATG5, ATG16L1, WIPI2, Beclin1, ATG14, STX17, SNAP29, VAMP8, VPS13B, VPS13C, VPS13D, ATG9A, ATG2A&B, TMEM41B, VMP1 or overexpressing ATG4B-C74S in the absence or presence of 500 nM Bafilomycin A1 for 1.5 h.

**e.** Quantification of the ratio of lipidated LC3 to tubulin with the control set as 1.00 (control cells without Bafilomycin A1) analyzed in (d) (mean ± SEM). Three experiments were performed for the statistics (two-tailed t-test). ***, $p<0.001$; ****, $p<0.0001$.

**f.** Immunofluorescence analysis the colocalization of GABARAP and LC3 in the KRAS(G12V)-expressing FHC cells.

**g.** Immunofluorescence of the mCherry-pHluorin-LC3B in the KRAS(G12V)-expressing 293T cells with control or knocking down of ULK1, FIP200, Beclin1, ATG14, WIPI2, ATG5, ATG16L1, ATG2A&B, VPS13B, VPS13C, VPS13D, STX17, SNAP29, VAMP8, ATG9A, TMEM41B, VMP1 or overexpressing ATG4B-C74S. Representative cell images are shown. Scale bar sizes are indicated in the image.

**h.** Quantification of the yellow ($RFP^+GFP^+$) and Red ($RFP^+GFP^-$) LC3 puncta. Data are represented as mean ± SEM. Three experiments (50 cells for each group/experiment) were performed for the statistics (two-tailed t-test). *, $p<0.05$.

**i.** Immunoblotting and in-gel fluorescence detection of in the KRAS(G12V) and control cells stably expressing HaloTag (Halo)-LC3B pulse-labeled for 20 min with 100 nM tetramethyl rhodamine (TMR)-conjugated ligand in nutrient-rich medium with control or knocking down of ULK1, FIP200, Beclin1, ATG14, WIPI2, ATG5, ATG16L1, ATG2A&B, VPS13B, VPS13C, VPS13D, STX17, SNAP29, VAMP8, ATG9A, TMEM41B, VMP1 or overexpressing ATG4B-C74S in the absence or presence of 500 nM Bafilomycin A1 for 1.5 h.

**j.** Quantification of results shown in (i). Halo-TMR band intensity was normalized by the sum of the band intensities Halo-TMR-LC3B and Halo-TMR, and the control set (KRAS(G12V) 293T cells with ligand without Bafilomycin A1) as 1.00. Three experiments were performed for the statistics (two-tailed t-test). ***, $p<0.001$; ****, $p<0.0001$.

**k.** FACS analysis of the control and KRAS(G12V) cells co-expressing mt-Keima and Parkin with control or knocking down of ULK1, FIP200, Beclin1, ATG14, WIPI2, ATG5, ATG16L1, ATG2A&B, VPS13B, VPS13C, VPS13D, STX17, SNAP29, VAMP8, ATG9A, TMEM41B, VMP1 or overexpressing ATG4B-C74S using V610 and Y610-mCherry detectors (Beckman CytoFLEX LX). The FACS results are representative of at least three independent experiments.

**l.** Quantification of results shown in (k). The percentage of cells with mitophagy based on Y610-mCherry/V610. Data are represented as mean ± SEM. Three experiments were performed for the statistics (two-tailed t-test). **, $p<0.01$; ***, $p<0.001$; ****, $p<0.0001$.

**m.** Immunoblot analysis of the LC3 lipidation from cells transfected with control or knocking down of Beclin1, ATG14, ATG9A, or ATG2A&B in starvation-induced autophagy by EBSS treatment in the absence or presence of 500 nM Bafilomycin A1 for 1.5 h.

**n.** Quantification of the ratio of lipidated LC3 to tubulin with the control set as 1.00 (KRAS(G12V) cells with Bafilomycin A1) analyzed in (m) (mean ± SEM). Three experiments were performed for the statistics (two-tailed t-test). **, p<0.01; ***, $p<0.001$; ****, $p<0.0001$.

**o.** Immunofluorescence of 293T cells expressing mCherry-pHluorin-LC3B with control or knocking down of Beclin1, ATG14, ATG9A and ATG2A&B in starvation-induced autophagy by EBSS treatment. Representative cell images are shown. Scale bar sizes are indicated in the image.

**p.** Quantification of the yellow ($RFP^{+}GFP^{+}$) and Red ($RFP^{+}GFP^{-}$) LC3 puncta. Data are represented as mean ± SEM. Three experiments (50 cells for each group/experiment) were performed for the statistics (two-tailed t-test). *, $p<0.05$, **, $p<0.01$.

**q.** Heatmap to show the changes of LC3 lipidation (the ratio of lipidated LC3 to tubulin with the control set as 1.00, related to Supplementary information, Fig. S1m, n) and autophagic flux by the tandem fluorescent LC3 system (AG and AL, the control set as 1.00, related to Supplementary information, Fig. S1o, p) in the control and EBSS group with knocking down Beclin1, ATG14, ATG2A&2B, or ATG9A, respectively. Color represents the $\log_2$ (fold) of counts per samples.

**r.** Electron microscopy images of APEX2-labeled LC3 and the autophagosomes in multiple cancer cell lines with RAS mutations. Scale bar sizes are indicated in the image

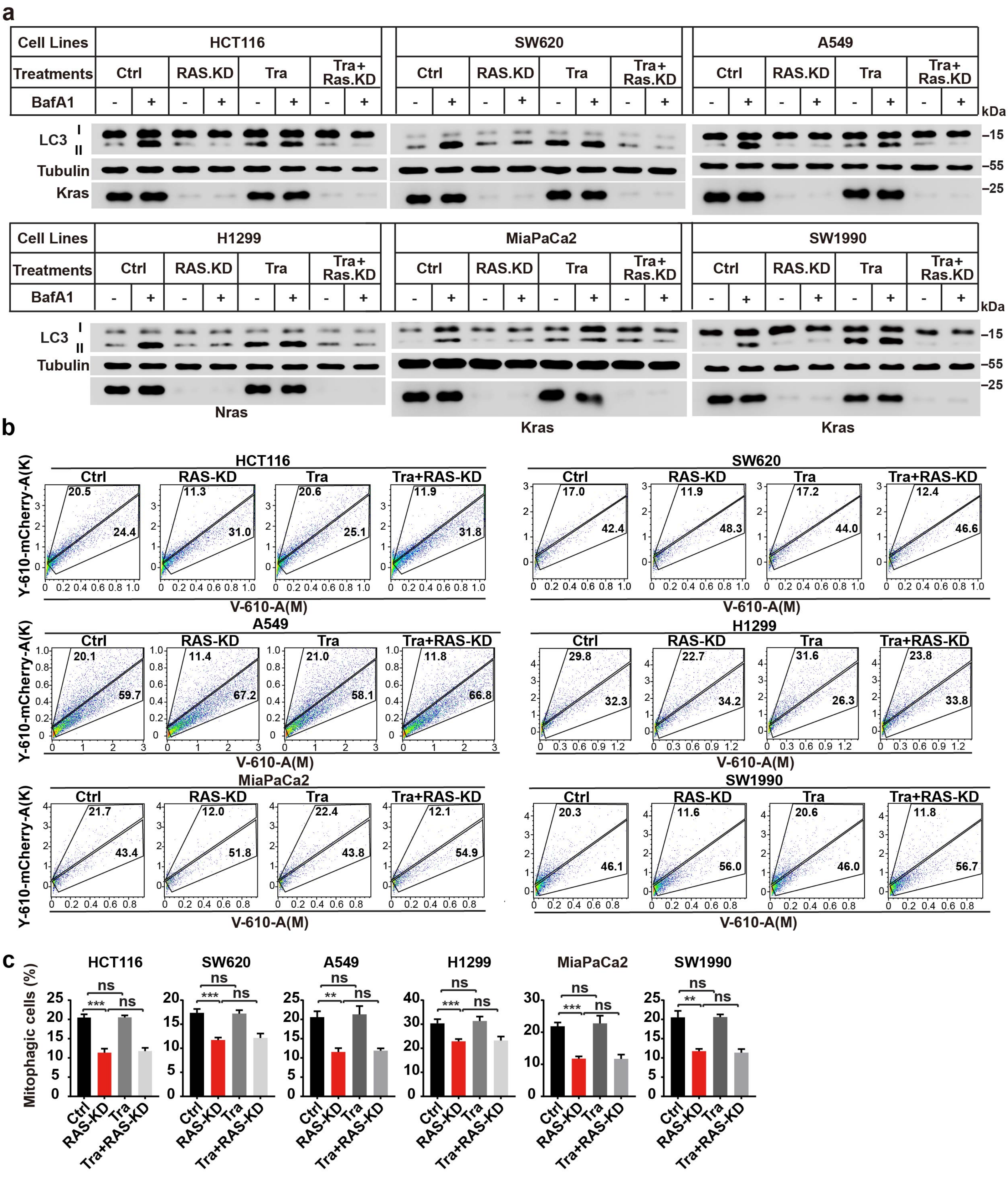

a
Cell Lines
Treatments
BafA1
HCT116
SW620
A549
H1299
MiaPaCa2
SW1990
Ctrl
RAS.KD
Tra
Tra+ Ras.KD
LC3
I
II
Tubulin
Kras
Nras
kDa
–15
–55
–25
b
Ctrl
RAS-KD
Tra
Tra+RAS-KD
Y-610-mCherry-A(K)
V-610-A(M)
c
Mitophagic cells (%)
ns
***
**

**Figure. S2 RAS knockdown led to decreased autophagosome biogenesis and autophagic flux**

**a.** Immunoblot analysis of LC3 lipidation of HCT116, SW620, A549, H1299, MiaPaCa-2 and SW1990 cells with or without RAS knocking down and Trametinib (Tra, 1 μM) treatment for 24 h in the absence or presence of 500 nM Bafilomycin A1 for 1.5 h.

**b.** FACS analysis of HCT116, SW620, A549, H1299, MiaPaCa-2 and SW1990 cells co-expressing mt-Keima and Parkin with or without RAS knocking down and Trametinib (Tra, 10 μM) treatment using V610 and Y610-mCherry detectors (Beckman CytoFLEX LX). The FACS results are representative of at least three independent experiments.

**c.** The percentage of cells with mitophagy based on Y610-mCherry/V610 calculated for (b). Data are represented as mean ± SEM. Three experiments were performed for the statistics (two-tailed t-test). **, $p<0.01$; ***, $p<0.001$.

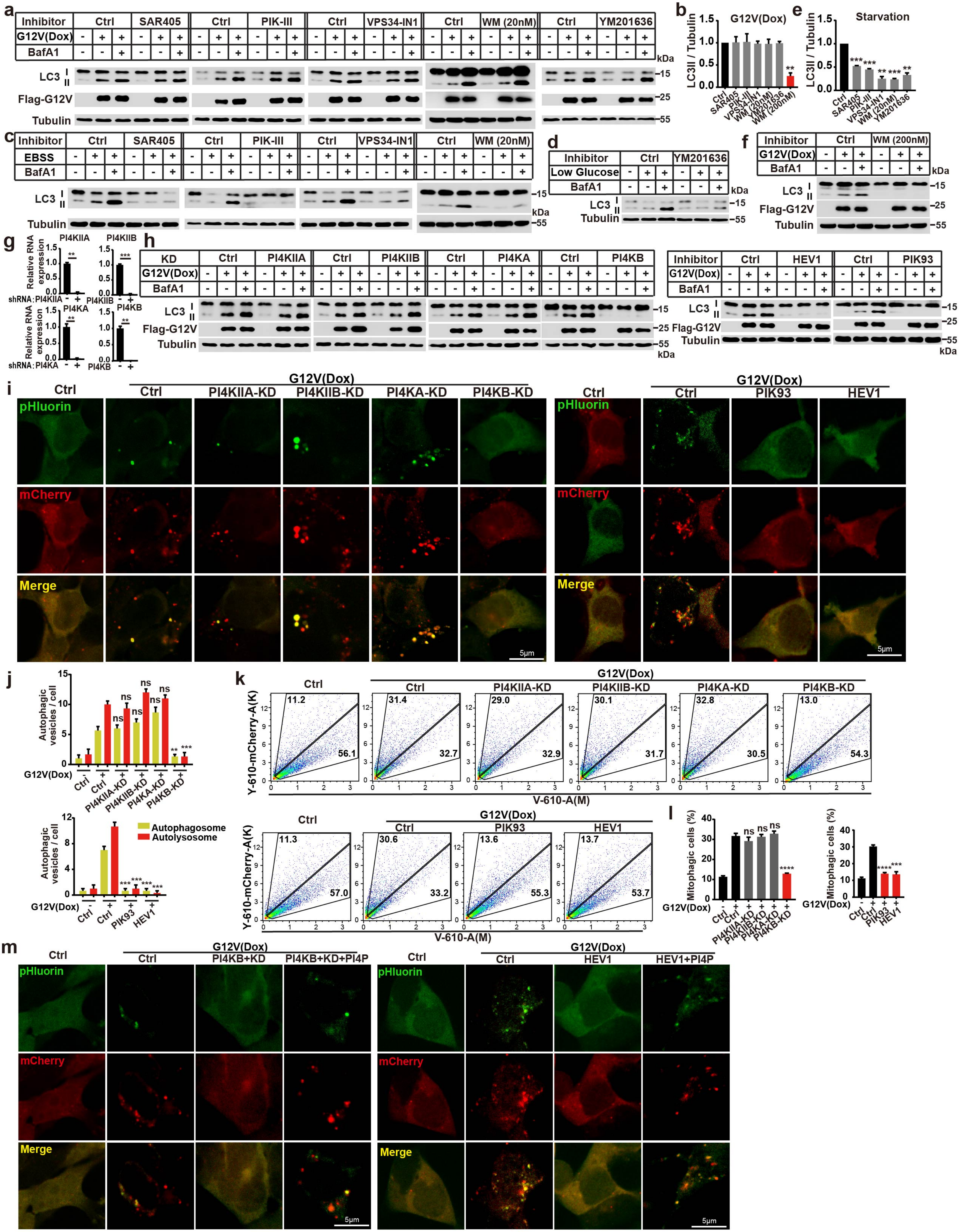
a
Inhibitor
Ctrl
SAR405
PIK-III
VPS34-IN1
WM (20nM)
YM201636
G12V(Dox)
BafA1
LC3
Flag-G12V
Tubulin
kDa
b
G12V(Dox)
LC3II / Tubulin
WM (200nM)
e
Starvation
c
EBSS
d
Low Glucose
f
WM (200nM)
g
PI4KIIA
PI4KIIB
PI4KA
PI4KB
Relative RNA expression
shRNA:
h
KD
HEV1
PIK93
i
pHluorin
mCherry
Merge
PI4KIIA-KD
PI4KIIB-KD
PI4KA-KD
PI4KB-KD
5μm
j
Autophagic vesicles / cell
Autophagosome
Autolysosome
k
Y-610-mCherry-A(K)
V-610-A(M)
l
Mitophagic cells (%)
m
PI4KB+KD
PI4KB+KD+PI4P
HEV1+PI4P

**Figure. S3 The PI3K and PI5K complexes do not regulate RINCAA, but regulate the starvation-induced autophagy**

**a.** Immunoblot analysis of LC3 lipidation in KRAS(G12V) and control cells treated with SAR405 (40 nM), PIK-III (40 nM), VPS34-IN1 (40 nM), Wortmannin (20 nM), or YM201636 (100 nM) in the absence or presence of 500 nM Bafilomycin A1 for 1.5 h.

**b.** Quantification of results shown in (a) (mean ± SEM). Three experiments were performed for the statistics (two-tailed t-test). **, $p<0.01$.

**c.** Immunoblot analysis of LC3 lipidation of cells treated with SAR405 (40 nM), PIK-III (40 nM), VPS34-IN1 (40 nM), or Wortmannin (20 nM) in starvation-induced autophagy by EBSS treatment in the absence or presence of 500 nM Bafilomycin A1 for 1.5 h.

**d.** Immunoblot analysis of LC3 lipidation of cells treated with YM201636 (100 nM) in low glucose-induced autophagy in the absence or presence of 500 nM Bafilomycin A1 for 1.5 h.

**e.** Quantification of results shown in (c, d) (mean ± SEM). Three experiments were performed for the statistics (two-tailed t-test). **, $p<0.01$; ***, $p<0.001$.

**f.** Immunoblot analysis of LC3 lipidation in KRAS(G12V) and control cells treated with Wortmannin (200 nM) in the absence or presence of 500 nM Bafilomycin A1 for 1.5h.

**g.** Relative RNA expression of cells transfected with control or shRNAs against PI4KIIA, PI4KIIB, PI4KA or PI4KB (mean ± SEM). Three experiments were performed for the statistics (two-tailed t-test). **, $p<0.01$; ***, $p<0.001$.

**h.** Immunoblot analysis of LC3 lipidation in KRAS(G12V) and control cells with Knocking down of PI4Ks or treatment of PI4KB inhibitors in the absence or presence of 500 nM Bafilomycin A1 for 1.5 h.

**i.** Immunofluorescence of KRAS(G12V) and control cells expressing mCherry-pHluorin-LC3B with Knocking down of PI4Ks or treatment of PI4KB inhibitors.

**j.** Quantification of the results in (i) (mean ± SEM). Three experiments (50 cells for each group/experiment) were performed for the statistics (two-tailed t-test). **, $p<0.01$; ***, $p<0.001$.

**k.** FACS analysis of control and KRAS(G12V) cells co-expressing mt-Keima and Parkin with control or knocking down of PI4Ks using V610 and Y610-mCherry detectors

(Beckman CytoFLEX LX). The FACS results are representative of at least three independent experiments.

**l.** Quantification of results shown in (k). The percentage of cells with mitophagy based on Y610-mCherry/V610. Data are represented as mean ± SEM. Three experiments were performed for the statistics (two-tailed t-test). ***, $p<0.001$; ****, $p<0.0001$.

**m.** Immunofluorescence of 293T cells expressing mCherry-pHluorin-LC3B were rescued by PI4P with suppressed PI4KB by knocking down or treatment of the inhibitor. Representative cell images are shown. Scale bar sizes are indicated in the image.

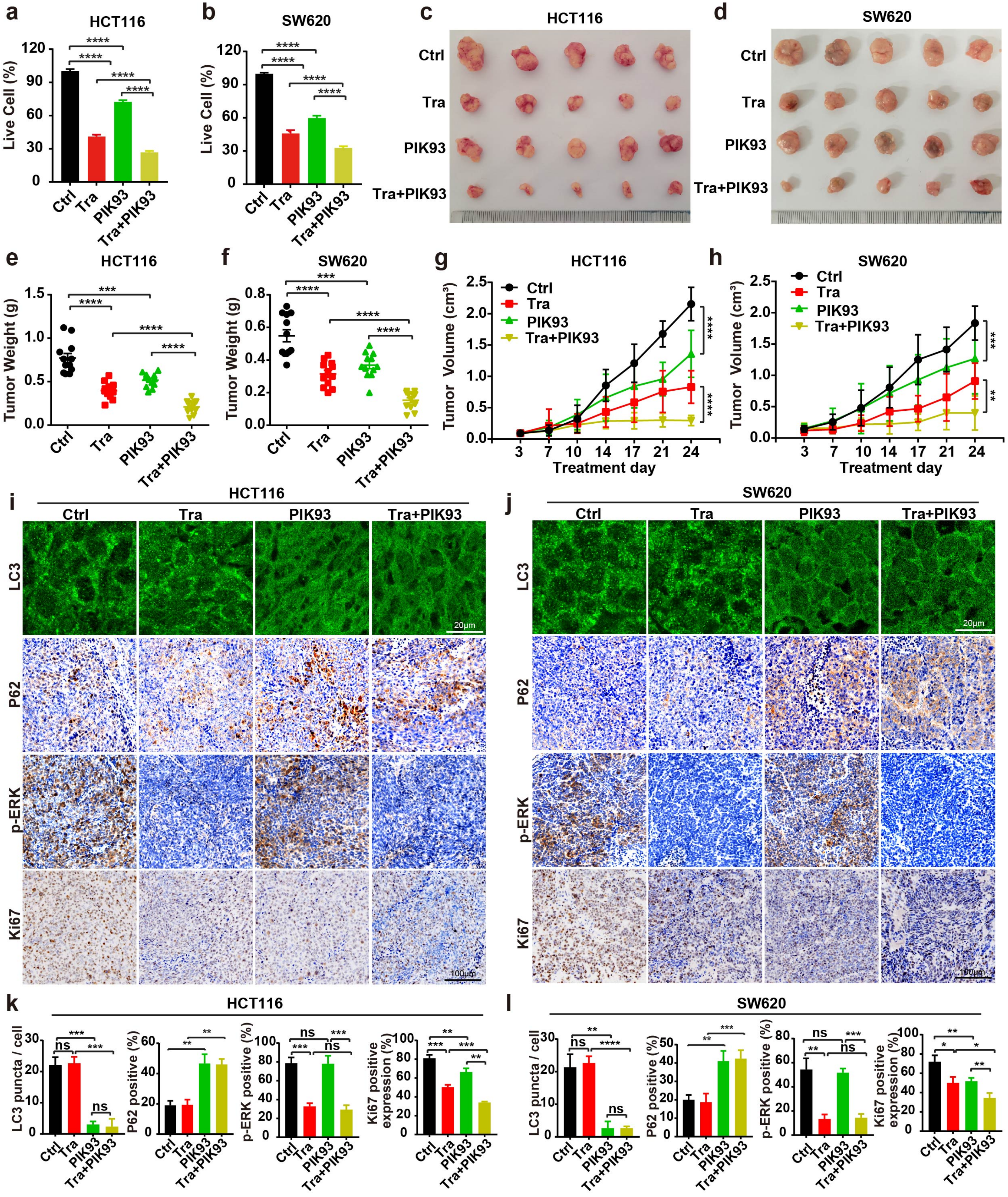
a
HCT116
Live Cell (%)
Ctrl
Tra
PIK93
Tra+PIK93
b
SW620
c
HCT116
d
SW620
e
Tumor Weight (g)
f
g
Tumor Volume (cm³)
Treatment day
h
i
j
LC3
P62
p-ERK
Ki67
20μm
100μm
k
l
LC3 puncta / cell
P62 positive (%)
p-ERK positive (%)
Ki67 positive expression (%)
ns

**Figure. S4 PI4KB inhibition slows xenograft tumors with RAS mutation**

**(a, b)** HCT116 and SW620 cells were treated with trametinib (100 nM) and PIK93 (1 μM) for 96 h, and the cell viability was analyzed (mean ± SEM). Three experiments were performed for the statistics (two-tailed t-test). ****, $p<0.0001$.

**(c, d)** Images of xenograft tumors of HCT116 and SW620 cells from mice treated with: (1) vehicle (Ctrl); (2) Trametinib (Tra); (3) PIK93; or (4) the combination of both (Tra+PIK93) (n = 12 mice (one tumor/mice) in each group). The tumors were removed and photographed after 24-day treatment.

**(e, f)** Weights of the xenograft tumors in (c, d) (means ± SD). The statistical analysis was performed by two-tailed t-test. *, $p<0.05$; **, $p<0.01$; ****, $p<0.0001$.

**(g, h)** The growth curve of the xenograft tumors in (c, d) (means ± SD). Statistical analysis was performed by two-way-ANOVA; ****, $p<0.0001$.

**(i, j)** Immunofluorescence and immunohistochemical analysis of the sections of xenograft tumors in (C, D). Sections were stained with antibody against LC3, p-ERK1/2 or Ki67, as indicated. Scale bar sizes are indicated in the image.

**(k, l)** Statistical analysis of the numbers of LC3 puncta, p-ERK positive rates, and levels of Ki67 expression in (I, J) (means ± SD). Statistical analysis was performed by two-tailed t-test; *, $p<0.05$; **, $p<0.01$; ***, $p<0.001$; ****, $p<0.0001$.

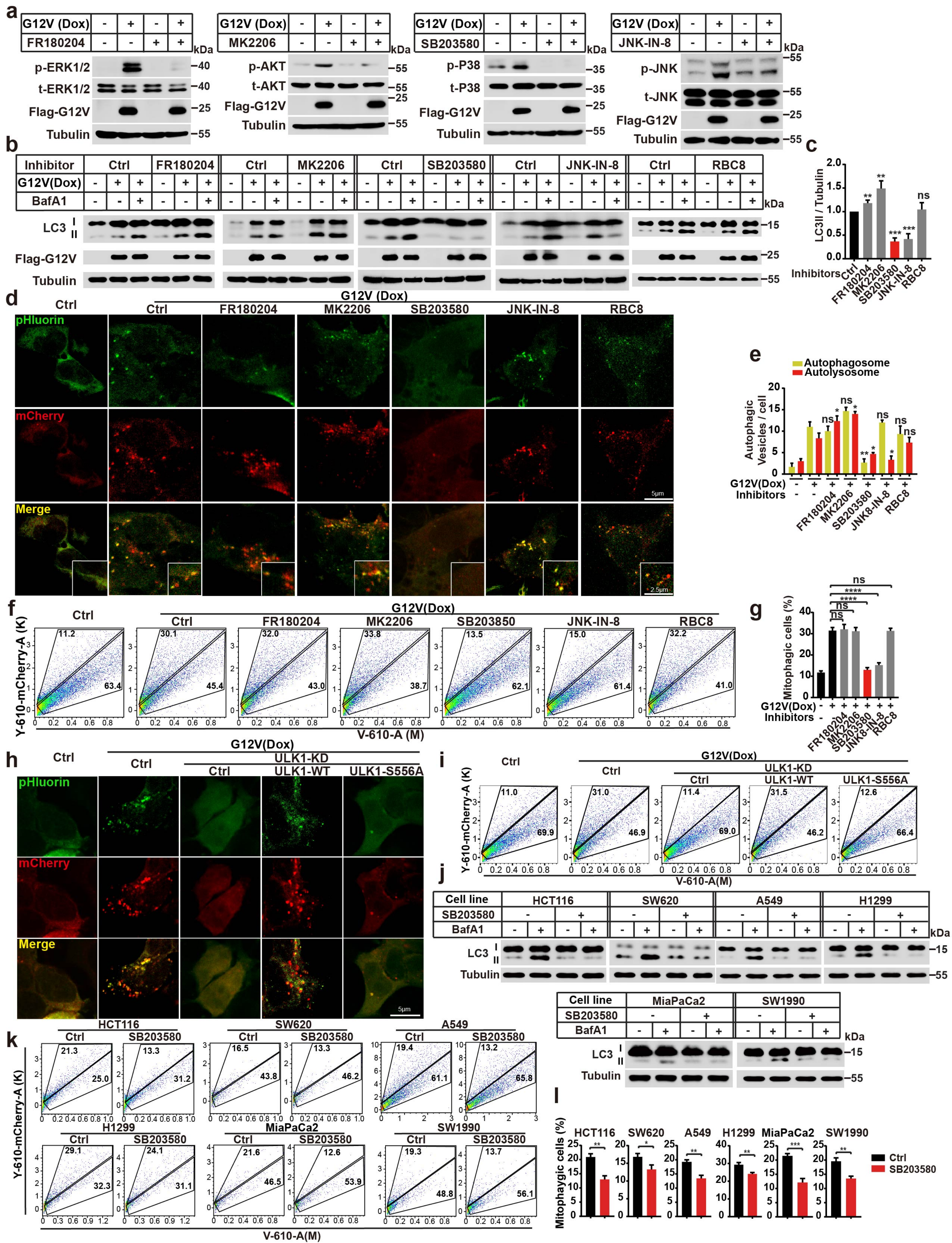
a
G12V (Dox)
FR180204
p-ERK1/2
t-ERK1/2
Flag-G12V
Tubulin
MK2206
p-AKT
t-AKT
SB203580
p-P38
t-P38
JNK-IN-8
p-JNK
t-JNK
kDa
b
Inhibitor
Ctrl
FR180204
MK2206
SB203580
JNK-IN-8
RBC8
G12V(Dox)
BafA1
LC3 I II
Flag-G12V
Tubulin
c
LC3II / Tubulin
Inhibitors
ns
d
Ctrl
G12V (Dox)
pHluorin
mCherry
Merge
5μm
2.5μm
e
Autophagosome
Autolysosome
Autophagic Vesicles / cell
G12V(Dox)
Inhibitors
JNK8-IN-8
f
G12V(Dox)
SB203850
Y-610-mCherry-A (K)
V-610-A (M)
g
Mitophagic cells (%)
h
ULK1-KD
ULK1-WT
ULK1-S556A
i
V-610-A(M)
j
Cell line
HCT116
SW620
A549
H1299
MiaPaCa2
SW1990
k
l
Ctrl
SB203580

**Figure. S5 The P38-pathway is required for RINCAA**

**a.** Immunoblot analysis of levels of p-ERK, p-AKT, p-P38 or p-JNK in control and KRAS(G12V)-cells treated with FR180204 (10 μM), MK2206 (10 μM), SB203580 (10 μM) or JNK-IN-8 (10 μM).

**b.** Immunoblot analysis of LC3 lipidation of the KRAS(G12V)-cells treated with FR180204 (10 μM), MK2206 (10 μM), SB203580 (10 μM), or RBC8 (10 μM) in the absence or presence of 500 nM Bafilomycin A1 for 1.5 h.

**c.** Quantification of the results in (b) (mean ± SEM). Three experiments were performed for the statistics (two-tailed t-test). **, $p$<0.01; ***, $p$<0.001.

**d.** Immunofluorescence of 293T cells expressing mCherry-pHluorin-LC3B treated with FR180204 (10 μM), MK2206 (10 μM), SB203580 (10 μM), or RBC8 (10 μM). Representative cell images are shown. Scale bar sizes are indicated in the image.

**e.** Quantification of the yellow ($RFP^+GFP^+$) and Red ($RFP^+GFP^-$) LC3 puncta in (d). Data are represented as mean ± SEM. Three experiments (50 cells for each group/experiment) were performed for the statistics (two-tailed t-test). *, p<0.05; **, p<0.01.

**f.** FACS analysis of control and KRAS(G12V) cells co-expressing mt-Keima and Parkin treated with FR180204 (10 μM), MK2206 (10 μM), SB203580 (10 μM), or RBC8 (10 μM) using V610 and Y610-mCherry detectors (Beckman CytoFLEX LX). The FACS results are representative of at least three independent experiments.

**g.** Quantification of results shown in (f). The percentage of cells with mitophagy based on Y610-mCherry/V610. Data are represented as mean ± SEM. Three experiments were performed for the statistics (two-tailed t-test). ****, $p$<0.0001.

**h.** Immunofluorescence of 293T cells expressing mCherry-pHluorin-LC3B treated with FR180204 (10 μM), MK2206 (10 μM), SB203580 (10 μM), or RBC8 (10 μM). Representative cell images are shown. Scale bar sizes are indicated in the image.

**i.** FACS analysis of control and KRAS(G12V) cells co-expressing mt-Keima and Parkin.To analyze the rescue effects of WT-ULK1 and ULK1-S556A on the KRAS(G12V)-induced mitophagy. The percentage of cells with mitophagy based on Y610-mCherry/V610. The FACS results are representative of at least three independent experiments.

**j.** Immunoblot analysis of LC3 lipidation of the different cancer cell lines treated with or without SB203580 (10 μM) in the absence or presence of 500 nM Bafilomycin A1 for 1.5 h.

**k.** FACS analysis of the different cancer cell lines co-expressing mt-Keima and Parkin treated with or without SB203580 (10 μM) The percentage of cells with mitophagy based on Y610-mCherry/V610. The FACS results are representative of at least three independent experiments.

**l.** Quantification of results shown in (k). The percentage of cells with mitophagy based on Y610-mCherry/V610. Data are represented as mean ± SEM. Three experiments were performed for the statistics (two-tailed t-test). ****, $p$<0.0001.

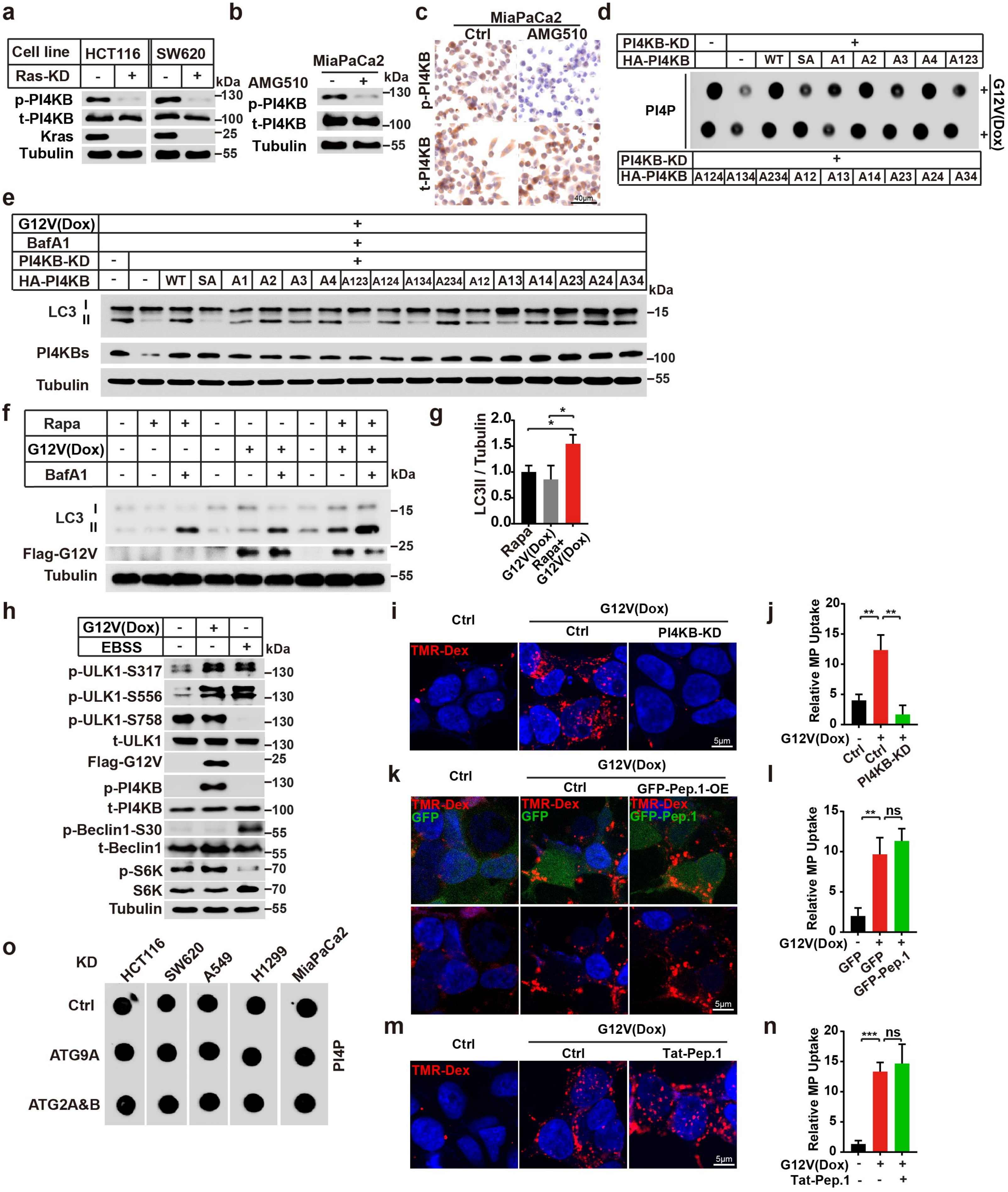

a
Cell line
HCT116
SW620
Ras-KD
p-PI4KB
t-PI4KB
Kras
Tubulin
kDa
b
MiaPaCa2
AMG510
p-PI4KB
t-PI4KB
Tubulin
c
MiaPaCa2
Ctrl
AMG510
p-PI4KB
t-PI4KB
40μm
d
PI4KB-KD
HA-PI4KB
WT SA A1 A2 A3 A4 A123
PI4P
G12V(Dox)
A124 A134 A234 A12 A13 A14 A23 A24 A34
e
G12V(Dox)
BafA1
PI4KB-KD
HA-PI4KB
LC3
PI4KBs
Tubulin
f
Rapa
G12V(Dox)
BafA1
LC3
Flag-G12V
Tubulin
g
LC3II / Tubulin
Rapa
G12V(Dox)
Rapa+ G12V(Dox)
h
G12V(Dox)
EBSS
p-ULK1-S317
p-ULK1-S556
p-ULK1-S758
t-ULK1
Flag-G12V
p-PI4KB
t-PI4KB
p-Beclin1-S30
t-Beclin1
p-S6K
S6K
Tubulin
i
Ctrl
G12V(Dox)
Ctrl
PI4KB-KD
TMR-Dex
5μm
j
Relative MP Uptake
G12V(Dox)
Ctrl
Ctrl
PI4KB-KD
k
Ctrl
G12V(Dox)
Ctrl
GFP-Pep.1-OE
TMR-Dex
GFP
GFP-Pep.1
5μm
l
Relative MP Uptake
G12V(Dox)
GFP
GFP
GFP-Pep.1
m
Ctrl
G12V(Dox)
Ctrl
Tat-Pep.1
TMR-Dex
5μm
n
Relative MP Uptake
G12V(Dox)
Tat-Pep.1
o
KD
HCT116
SW620
A549
H1299
MiaPaCa2
Ctrl
ATG9A
ATG2A&B
PI4P

**Figure. S6 Validation of the PI4KB-peptide 1 phosphorylation antibody, and pinpointing the phosphorylation of PI4KB**

**a.** Immunoblot analysis of the PI4KB phosphorylation by the antibody against p-PI4KB in HCT116 and SW620 cells with or without RAS knocking down.

**b.** Immunoblot analysis of the PI4KB phosphorylation by the antibody against p-PI4KB in MiaPaCa-2 cells treated with or without with 10 μM AMG510 for 1.5 h.

**c.** Immunohistochemical analysis of the sections of MiaPaCa-2 cells treated with or without with 10 μM AMG510 for 1.5 h. Sections were stained with antibody against PI4KB and p-PI4KB as indicated. Scale bar sizes are indicated in the image.

**d.** Dot blot analysis of the effect of PI4KB mutants on the KRAS(G12V)-induced PI4P generation in 293T cells.

**e.** Immunoblot analysis of the rescue effect of PI4KB-WT, PI4KB-SA and PI4KB mutants on KRAS(G12V)-induced LC3 lipidation after knocking down PI4KB in 293T cells in the absence or presence of 500 nM Bafilomycin A1 for 1.5 h.

**f.** Immunoblot analysis of the LC3 lipidation of the cell lysates from the control or KRAS(G12V) cells treated with 2 μM Rapamycin for 1.5 h in the absence or presence of 500 nM Bafilomycin A1 for 1.5 h.

**g.** Quantification of the results in (f) (mean ± SEM). Three experiments were performed for the statistics (two-tailed t-test). *, $p$<0.05.

**h.** Immunoblot analysis of the changes of the different sites of p-ULK, the level of p-PI4KB, p-Beclin1 and p-S6K in the KRAS(G12V) and starvation 293T cells.

**i.** Macropinocytosis visualization using TMR-Dex in control and KRAS(G12V) 293T cells with or without PI4KB knocking down. Representative cell images are shown. Scale bar sizes are indicated in the image.

**j.** Quantification of macropinocytosis in (I). Data are represented as mean ± SEM. Three experiments (50 cells for each group/experiment) were performed for the statistics (two-tailed t-test). **, p<0.01.

**k.** Macropinocytosis visualization using TMR-Dex in control and KRAS(G12V) 293T cells with or without GFP-Peptide1 (GFP-Pep.1) overexpression. Representative cell images are shown. Scale bar sizes are indicated in the image.

**l.** Quantification of macropinocytosis in (k). Data are represented as mean ± SEM. Three experiments (50 cells for each group/experiment) were performed for the statistics (two-tailed t-test). **, $p<0.01$.

**m.** Macropinocytosis visualization using TMR-Dex (1 mg/mL) in control and KRAS(G12V) 293T cells with or without Tat-Peptide1 (Tat-Pep.1, 25 μM) treatment for 2 h. Representative cell images are shown. Scale bar sizes are indicated in the image.

**n.** Quantification of macropinocytosis in (m). Data are represented as mean ± SEM. Three experiments (50 cells for each group/experiment) were performed for the statistics (two-tailed t-test). ***, $p<0.001$.

**Supplementary information, Table S1 MS of p-PI4KB**

| Name | Sequence | Modifications | % of phosphated peptides | | |
|---|---|---|---|---|---|
| | | | Con | ULK | ULK+IN |
| Peptide 1 | KRELPSLSPAPDTGLSPSK | S6(Phospho); S14(Phospho) | 0.35 | 0.61 | 0.14 |
| Peptide 2 | SKSDATASISLSSNLKR | S3(Phospho) | 0.55 | 0.45 | 0.46 |
| Peptide 3 | RTAsNPKVENEDEPVR | T2(Phospho); S4(Phospho) | 1.00 | 1.00 | 1.00 |
| Peptide 4 | STRsVENLPEcGITHEQR | S1(Phospho); S4(Phospho) | 1.00 | 1.00 | 1.00 |
| Peptide 5 | RLSEQLAHTPTAFKR | S2(Phospho); S3(Phospho); S4(Phospho) | 0.48 | 0.61 | 0.77 |

**Supplementary information, Table S2 Antibodies used in this study**

| Antibodies | Source | Identifier |
|---|---|---|
| Mouse monoclonal anti-FLAG | Sigma | Cat# F3165; RRID: AB_259529 |
| Rabbit monoclonal anti-FLAG | CST | Cat# 14793; RRID: AB_2572291 |
| Rat monoclonal anti-FLAG | Novus | Cat# NBP1-06712SS; RRID: AB_1625982 |
| Rabbit monoclonal anti-HA | CST | Cat# 3724; RRID: AB_1549585 |
| Mouse monoclonal anti-Myc | CST | Cat# 2276; RRID: AB_331783 |
| Rabbit polyclonal anti-LC3 (WB) | ABclonal | Cat# A19665; RRID: AB_2862723 |
| Rabbit polyclonal anti-LC3 (IF) | MBL | Cat# PM036; RRID: AB_2274121 |
| Mouse monoclonal anti-LC3 (IF) | MBL | Cat# M152-3; RRID: AB_1279144 |
| Rabbit polyclonal anti-LC3 (IHC) | ABcepta | Cat# AP1802a, RRID: AB_2137695 |
| Mouse monoclonal anti-β-Tubulin | Zen Bioscience | Cat# 200608; RRID: AB_2722706 |
| Rabbit monoclonal anti-ULK1 | CST | Cat# 8054; RRID: AB_11178668 |
| Rabbit monoclonal anti-Phospho-ULK1 (Ser555) | CST | Cat#5869; RRID: AB_10707365 |
| Rabbit polyclonal anti-ULK1 (Ser556) conjugated to Biotin | Biorbyt | Cat# orb502261 |
| Rabbit monoclonal anti-Phospho-ULK1 (Ser317) | CST | Cat#6887S; RRID: AB_10831845 |
| Rabbit monoclonal anti-Phospho-ULK1 (Ser757) | CST | Cat#14202S; RRID: AB_2665508 |
| Rabbit monoclonal anti-Phospho-Becline-1(Ser30) | CST | Cat#35955S; |
| Rabbit polyclonal anti-RB1CC1(FIP200) | Proteintech | Cat# 17250-1-AP; RRID: AB_10666428 |
| Mouse monoclonal anti-ATG5 | MBL | Cat# M153-3; RRID: AB_1278760 |
| Mouse monoclonal anti-ATG16L1 | MBL | Cat# M150-3; RRID: AB_1278758 |
| Mouse monoclonal anti-WIPI2 | Bio-Rad | Cat# MCA5780GA; RRID: AB_10845951 |
| Rabbit polyclonal anti-mCherry | This paper | N/A |
| Rabbit polyclonal anti-Beclin1 | Sigma | Cat# PRS3613; RRID: AB_1845329 |
| Rabbit polyclonal anti-ATG14 | MBL | Cat# PD026; |

| | | |
|---|---|---|
| | | RRID: AB_1953054 |
| Rabbit polyclonal anti-ATG9A | MBL | Cat# PD042; RRID: AB_2714019 |
| Rabbit monoclonal anti-ATG9A | CST | Cat# 13509; RRID: AB_2798241 |
| Rabbit polyclonal anti-ATG2A | MBL | Cat# PD041; RRID: AB_2810871 |
| Rabbit polyclonal anti-ATG2B | Sigma | Cat# HPA019665; RRID: AB_2274278 |
| Mouse anti-PI4P | Echelon | Cat# Z-P004; RRID: AB_11127796 |
| Rabbit polyclonal anti-PI4KB | Abcam | Cat# Ab134756 |
| Rabbit monoclonal anti-GFP (WB) | CST | Cat# 2956; RRID: AB_1196615 |
| Rabbit monoclonal anti-ERK | CST | Cat# 4695; RRID: AB_390779 |
| Rabbit monoclonal anti-Phospho-ERK | CST | Cat# 4370; RRID: AB_2315112 |
| Rabbit monoclonal anti-AKT | CST | Cat# 4691; RRID: AB_915783 |
| Rabbit monoclonal anti-Phospho-AKT | CST | Cat# 4060; RRID: AB_2315049 |
| Rabbit monoclonal anti-P38 | CST | Cat# 8690; RRID: AB_10999090 |
| Rabbit monoclonal anti-Phospho-P38 | CST | Cat# 4511; RRID: AB_213968 |
| Rabbit polyclonal anti-JNK | CST | Cat# 9252; RRID: AB_2250373 |
| Rabbit polyclonal anti-Phospho-JNK | CST | Cat# 4668; RRID: AB_823588 |
| Rabbit polyclonal anti-Ki67 | Abcam | Cat# ab15580, RRID: AB_443209 |
| Rabbit polyclonal anti-Ribophorin1(RPN1) | Dr. Randy Schekman | N/A |
| Rabbit polyclonal anti-Phospho-PI4KB | N/A | N/A |
| Rabbit polyclonal anti-STX17 | Proteintech | Cat# 17815-1-AP, RRID: AB_2255542 |
| Rabbit polyclonal anti-SNAP29 | Proteintech | Cat# 12704-1-AP, RRID: AB_2192340 |
| Rabbit polyclonal anti-VAMP8 | Proteintech | Cat# 15546-1-AP, RRID: AB_2878150 |
| Rabbit polyclonal anti-KRAS | ABclonal | Cat# A1190, RRID: AB_2758846 |
| Rabbit polyclonal anti-NRAS | ABclonal | Cat# A7566 RRID: AB_2770660 |

| | | |
|---|---|---|
| Rabbit polyclonal anti-p70 S6K | CST | Cat# 2708S<br>RRID: AB_390722 |
| Rabbit polyclonal anti-p-p70 S6K | CST | Cat# 9025S<br>RRID: AB_2734746 |

**Supplementary information, Table S3 Oligonucleotides used in this study**

| | |
|---|---|
| **Oligos for siRNAs** | |
| ULK1 siRNA target sequence-1: | CGCGCGGTACCTCCAGAGCAA |
| ULK1 siRNA target sequence-2: | TGCCCTTTGCGTTATATTGTA |
| FIP200 siRNA target sequence-1: | CTGGGACGGATACAAATCCAA |
| FIP200 siRNA target sequence-2: | ACGCAAATCAGTTGATGATTA |
| ATG5 siRNA target sequence-1: | AACCTTTGGCCTAAGAAGAAA |
| ATG5 siRNA target sequence-2: | CTAGGAGATCTCCTCAAAGAA |
| ATG5 siRNA target sequence-3: | AAGACTTACCGGACCACTGAA |
| ATG5 siRNA target sequence-4: | CATCATAGCTTTATTACTCTA |
| ATG16L1 siRNA target sequence-1: | GGAGATCATCCTGCAGTATAA |
| ATG16L1 siRNA target sequence-2: | CACGAGATAAGTCCCGGACAT |
| ATG16L1 siRNA target sequence-3: | CTCCCGTGATGACTTGCTAAA |
| ATG16L1 siRNA target sequence-4: | CAGGATCCAGTTGCAATGATA |
| WIPI2 siRNA target sequence-1: | ACACGATTCTGGCTGTGAA |
| WIPI2 siRNA target sequence-2: | TGACGCAAGTGGAACTAAA |
| WIPI2 siRNA target sequence-3: | CGCTGTCAATCAACAACGA |
| Beclin1 siRNA target sequence-1: | GAGGATGACAGTGAACAGTTA |
| Beclin1 siRNA target sequence-2: | TGGACAGTTTGGCACAATCAA |
| Beclin1 siRNA target sequence-3: | AGGGTCTAAGACGTCCAACAA |
| Beclin1 siRNA target sequence-4: | ACCGACTTGTTCCTTACGGAA |
| ATG14 siRNA target sequence-1: | CTCGGTGACCTCCTGGTTTAA |
| ATG14 siRNA target sequence-2: | TTGGATTAGCCTCCCTAACAA |
| ATG14 siRNA target sequence-3: | CTGCATACCCTCAGGAATCTA |
| ATG14 siRNA target sequence-4: | CCGGGAGAGGTTTATCGACAA |
| ATG2A siRNA target sequence-1: | Dr, M. Zhang |
| ATG2B siRNA target sequence-2: | Dr, M. Zhang |
| PI4KB siRNA target sequence-1: | GCGACATGTTCAACTACTA |
| PI4KB siRNA target sequence-2: | GCACCATTCGAAACCTCAA |
| PI4KB siRNA target sequence-3: | CTTGCTCGATTACTTCCTA |
| **Oligos for shRNAs** | |
| Ctrl shRNA target sequence: | CAACAAGATGAAGAGCACCAA |
| ULK1shRNA target sequence-1: | GCCCTGGATACGTCTTGTAAT |
| ULK1shRNA target sequence-2: | GCCCTTTGCGTTATATTGTAT |
| PI4KIIA shRNA target sequence-1: | CCCTAACTTCGTCAAGGACTT |
| PI4KIIA shRNA target sequence-2: | CCTCTTCCTGAGAACACTAAC |
| PI4KIIB shRNA target sequence-1: | GACATGAACTTTGTGCAAGAT |
| PI4KIIB shRNA target sequence-2: | CCTGATGAATGGAGAGCATAT |
| PI4KA shRNA target sequence-1: | ACGACATGATCCAGTACTATC |
| PI4KA shRNA target sequence-2: | CAAGGCTGGATCAACACATAC |
| PI4KB shRNA target sequence-1: | GCAAGAAACACGAAGGATCAT |

| | |
|---|---|
| PI4KB shRNA target sequence-2: | CCACAGGCCATCCTCTTATT |
| VPS13B shRNA target sequence-1: | GCCTTACTCAACCTTCTGATA |
| VPS13B shRNA target sequence-2: | ATGGTACTCGCGCAGAATTTA |
| VPS13C shRNA target sequence-1: | CGGACAAAGTTAATCCAACAA |
| VPS13C shRNA target sequence-2: | GCAGTAACTAATGCCCTAAAT |
| VPS13D shRNA target sequence-1: | CCCAGAAAGTTCCAGATCAAA |
| VPS13D shRNA target sequence-2: | GCCAGATTTGACTTCAAGAAA |
| STX17 shRNA target sequence-1: | CGATCCAATATCCGAGAAATT |
| STX17 shRNA target sequence-2: | TTGTTATCAGATAGCGAAATC |
| SNAP29 shRNA target sequence-1: | CCAGAAACACATCAATAGCAT |
| SNAP29 shRNA target sequence-2: | ACAACCAAAGTGGACAAGTTA |
| VAMP8 shRNA target sequence-1: | CTTGGAACATCTCCGCAACAA |
| VAMP8 shRNA target sequence-2: | GATTGTCCTTATCTGCGTGAT |
| TMEM41B shRNA target sequence-1: | GTTGAACGTCATAGAGAACAT |
| TMEM41B shRNA target sequence-2: | GTTTCCTGGAACTCAATATTT |
| VMP1 shRNA target sequence-1: | GCAATGAACAAGGAACATCAT |
| VMP1 shRNA target sequence-2: | CGATGCAATCCACCTTGTGTT |
| **Oligos for Primers** | |
| PI4KIIA forward primer: | TTGTCCTTAACCAGGGCTATCT |
| PI4KIIA reverse primer: | GTACGGGGAACAATGTTGAGT |
| PI4KIIB forward primer: | ACCCAAATCAGAAGAGCCTTATG |
| PI4KIIB reverse primer: | CAAGGGCAGCAGACCTTATG |
| PI4KA forward primer: | AACCTGGACATAACTGTCGGC |
| PI4KA reverse primer: | GAGCCCCGGTTGGTTTTCTT |
| PI4KB forward primer: | CCTGCTCAACCATAAGCTCCC |
| PI4KB reverse primer: | AGTTTTCTACGGACCTCGTACT |
| ACTIN forward primer: | CATGTACGTTGCTATCCAGGC |
| ACTIN reverse primer: | CTCCTTAATGTCACGCACGAT |
| VPS13B forward primer: | ATCGTCCTTTCCGTCAATATCAC |
| VPS13B reverse primer: | CAGCACCAAATCAGTTGCAGA |
| VPS13C forward primer: | ACTTGATAGTCTTAGCGCCTACT |
| VPS13C reverse primer: | ATCCAGTTTGGGCGTTTTGAG |
| VPS13D forward primer: | TACCGCCTCCGTAGTTACAAG |
| VPS13D reverse primer: | GTAAAGTGCAATCGACATCCCA |
| TMEM41B forward primer: | CAGAGTCGCCGAACGATCG |
| TMEM41B reverse primer: | CCTAGAGCCTTGGCATCATCCA |
| VMP1 forward primer: | GACCAGAGACGTGTAGCAATG |
| VMP1 reverse primer: | ACAATGCTTTGACGATGCCATAA |
